# An ESCRT grommet cooperates with a diffusion barrier to maintain nuclear integrity

**DOI:** 10.1101/2022.12.12.520126

**Authors:** Nicholas R. Ader, Linda Chen, Ivan V. Surovtsev, William L. Chadwick, Elisa C. Rodriguez, Megan C. King, C. Patrick Lusk

**Affiliations:** Department of Cell Biology, Yale School of Medicine, _295 Congress Ave_, New Haven, CT, 06520; Department of Physics, Yale University, New Haven, CT, 06511; Department of Molecular, Cell and Developmental Biology, Yale University, New Haven, CT, 06511

## Abstract

The molecular mechanisms by which the endosomal sorting complexes required for transport (ESCRT) proteins contribute to the integrity of the nuclear envelope (NE) barrier are not fully defined. Here, we leveraged the single NE hole generated by mitotic extrusion of the *Schizosaccharomyces pombe* spindle pole body (SPB) to reveal two modes of ESCRT function executed by distinct complements of ESCRT-III proteins, both depending on CHMP7/Cmp7. A grommet-like function is required to restrict the NE hole in anaphase B, while replacement of Cmp7 by a sealing module ultimately closes the NE in interphase. Without Cmp7, nucleocytoplasmic compartmentalization surprisingly remains intact despite NE discontinuities up to 550 nm, suggesting mechanisms to prevent diffusion through these holes. We implicate SPB proteins as key components of a diffusion barrier acting with Cmp7 in anaphase B. Thus, NE remodeling mechanisms cooperate with proteinaceous diffusion barriers beyond nuclear pore complexes to protect the nuclear compartment.

## INTRODUCTION

In eukaryotic cells, the genome is enclosed by the nuclear envelope (NE), which establishes a protected nuclear compartment. The NE is contiguous with the endoplasmic reticulum (ER) but is functionally specialized due to a unique proteome that populates its constituent inner, outer, and nuclear pore membranes. Nucleocytoplasmic compartmentalization relies on the integrity of these membranes and the selective barrier properties of nuclear pore complexes (NPCs) embedded in the NE (Dultz et al., 2022). During mitosis, the NE barrier presents a challenge for the spindle as it must access the duplicated chromosomes to faithfully segregate the genome to each daughter cell. Evolution has provided several solutions to this problem with a continuum of “closed” to “open” mitosis mechanisms employed in different organisms that reflects the degree to which the NE remains globally intact during chromosome segregation (Makarova and Oliferenko, 2016). In all scenarios, elaborate changes to the morphology of the NE during this period are universally observed, helping to ensure either the maintenance (in the case of a closed mitosis) or re-establishment (in the case of an open mitosis) of nucleocytoplasmic compartmentalization (Ungricht and Kutay, 2017).

These NE remodeling mechanisms are perhaps most elaborate at the end of the open mitosis of mammalian cells, when decondensing chromosomes are surrounded by nascent NE. Completion of NE sealing requires the closing of numerous NE fenestrations through an annular fusion mechanism thought to be executed by the endosomal sorting complexes required for transport (ESCRT) machinery, specifically the ESCRT-III proteins (Gu et al., 2017; Olmos et al., 2015; Olmos et al., 2016; Ventimiglia et al., 2018; Vietri et al., 2015; von Appen et al., 2020). A similar mechanism has been invoked in the sealing of nuclear pores housing defective NPCs (Thaller et al., 2019; Thaller et al., 2021; Webster et al., 2014; Webster et al., 2016) and at NE ruptures frequently observed in cancer cells or cells migrating through constrictions (Denais et al., 2016; Raab et al., 2016). While the events that underlie ESCRT-III mediated NE sealing have not been mechanistically delineated, parallels have been drawn with more extensively studied ESCRT-mediated membrane fission reactions. For example, during intralumenal vesicle (ILV) formation on endosomes, ESCRT-I/II recruit and activate the polymerization of ESCRT-III proteins (Vietri et al., 2020). ESCRT-III proteins can form single-and double-stranded homo and heteropolymer filaments *in vitro*; such filaments have affinity for negative or positive membrane curvature, perhaps influenced by the specific ESCRT-III subunit composition (McCullough et al., 2018; Pfitzner et al., 2021; Remec Pavlin and Hurley, 2020). The AAA+ ATPase VPS4 is required for the dynamic exchange of filament subunits with a monomer pool (Adell *et al*., 2017; Mierzwa et al., 2017) and for the ultimate membrane fission step (McCullough *et al*., 2018).

Despite the topological similarity between a NE hole and the neck of forming ILVs, prior studies reveal distinct ESCRT requirements at the NE that hint at potentially unique membrane remodeling and fission mechanisms. For example, ESCRT-I/II are dispensable for the recruitment of ESCRT-III to the NE (Webster *et al*., 2014). Further, there is an ESCRT-II/III fusion protein, CHMP7 (Chm7 in *Saccharomyces cerevisiae* and Cmp7 in *Schizosaccharomyces pombe/Schizosaccharomyces japonicus*), with a unique function at the NE. Interestingly, unlike most ESCRT-mediated membrane remodeling events that rely on peripheral membrane proteins, the recruitment of CHMP7 to the NE requires a direct physical interaction with an integral inner nuclear membrane protein, LEM2 (Heh1/Src1 in *S.c.* and Heh1/Lem2 in *S.p./S.j.*) (Gu *et al*., 2017; Webster *et al*., 2016). *In vitro* (von Appen *et al*., 2020) and *in vivo* (Thaller *et al*., 2019) evidence further supports that LEM2 family members also activate CHMP7 polymerization. Moreover, LEM2 and CHMP7 may form a membrane-embedded co-polymer that lines NE holes like an O-ring, surrounding microtubules that remain attached to kinetochores (von Appen *et al*., 2020). Lastly, although ESCRT-III subunits including CHMP1A,B (Did2 in yeasts), CHMP2A,B (Vps2 in *S.c.* and Did4 in *S.p./S.j.*), CHMP3 (Vps24 in yeasts), CHMP4A,B (Snf7 in *S.c.* and Vps32 in *S.p/S.j.*), and IST1 (Ist1 in yeasts) have been shown to be recruited to the NE alongside additional regulatory factors in metazoans including CC2D1B (Ventimiglia *et al*., 2018), Nesprin-2G (Wallis et al., 2021) and UFD1 (Olmos *et al*., 2015), we lack a quantitative spatiotemporal framework including the order, co-dependency, and copy number of ESCRT proteins at a NE sealing site. This information is critical to be able to fully elucidate how (and indeed whether) ESCRTs seal the NE and will also lay the groundwork for further delineation of shared and unique mechanisms that ESCRT proteins engage in to support a diverse range of membrane remodeling activities.

Despite clear evidence for recruitment of ESCRT proteins to the NE in late mitosis (Gu *et al*., 2017; Olmos *et al*., 2015; Olmos *et al*., 2016; Ventimiglia *et al*., 2018; Vietri *et al*., 2015; von Appen *et al*., 2020), several observations call into question whether the ESCRT proteins are indeed necessary and/or sufficient to seal the NE. First, compromising ESCRT by depletion of various subunits only delays, but does not abrogate, the re-establishment of the accumulation of a soluble nuclear import reporter in the nucleus after mitotic exit in human cells (Olmos *et al*., 2015; Olmos *et al*., 2016; von Appen *et al*., 2020) and the semi-open mitosis of *S. japonicus* (Lee et al., 2020; Pieper et al., 2020). This result, which was mirrored by orthogonal studies modeling the repair of interphase nuclear ruptures induced by laser ablation (Halfmann et al., 2019) or migration through constrictions (Denais *et al*., 2016; Raab *et al*., 2016), suggest that there are alternative, ESCRT-independent mechanisms by which nuclear compartmentalization is reestablished. Indeed, Chm7/Cmp7 is dispensable for cell viability and global nucleocytoplasmic compartmentalization in *S. cerevisiae* (Webster *et al*., 2016) and *S. pombe* (Dey et al., 2020; Gu *et al*., 2017). One attractive explanation for these observations is the existence of a redundant, ESCRT-independent NE sealing mechanism(s). Indeed, genetic evidence in *Caenorhabditis elegans* and *S. japonicus* suggests that altering phospholipid biosynthesis and/or membrane fluidity can mitigate the ill effects of a compromised ESCRT-dependent NE sealing pathway (Lee *et al*., 2020; Penfield et al., 2020). Compensation by an orthogonal and CHMP7/Cmp7-independent ESCRT-III sealing mechanism has also been proposed (Lee *et al*., 2020). It is worth considering, however, that most of these prior studies have relied on an indirect tool to assess the integrity of the NE, namely the nuclear accumulation of a soluble reporter containing a nuclear localization signal (NLS). Direct visualization of the continuity of the NE membranes themselves, although more challenging, is ultimately essential to address whether ESCRTs are necessary for NE sealing.

Here, we determine mechanisms of ESCRT function at the NE by taking advantage of the fission yeast *S. pombe*. *S. pombe* inserts and extrudes its spindle pole body (SPB) into and out of the NE at every mitotic entry and exit, respectively (Ding et al., 1997). SPB extrusion thus leaves a single NE hole that must be sealed in each daughter cell in a temporal and topological context similar, but much better defined than, the numerous NE fenestrations present at the final stages of metazoan NE reformation. We performed a comprehensive analysis of the complement, copy number, and relative timing of ESCRT recruitment to the SPB extrusion site correlated to an ultrastructural timeline of NE hole dynamics. In addition to providing definitive data that ESCRTs are essential for ultimately sealing the NE, we also reveal an unexpected role for ESCRTs in restricting the size of the NE hole during anaphase B in a manner akin to the function of a grommet. Last, we uncover a mechanism that can explain the paradox of an apparently intact nuclear compartment containing persistent NE holes: the existence of a compensatory proteinaceous diffusion barrier that is sensitive to perturbation with 1,6 hexanediol and depends on components of the SPB, including pericentrin/Pcp1. Our findings thus support both sealing dependent and independent functions for NE-recruited ESCRTs while providing evidence for additional diffusion barriers beyond NPCs that support the integrity of nucleocytoplasmic compartmentalization.

## RESULTS

### ESCRTs are recruited in temporally distinct waves to the SPB extrusion site

We hypothesized that Cmp7 and member(s) of the ESCRT-III family would be recruited to the single NE hole generated in each fission yeast daughter cell nucleus upon SPB extrusion during anaphase B (Ding *et al*., 1997) (Figure 1A). We therefore tested the localization of functional (see Methods) GFP-tagged ESCRT proteins, including: Cmp7, Vps32, Ist1, Vps24, Did4, and Did2 (Figure 1B, Supplementary Figure S1A, and Supplementary Table S1) in cells co-expressing Pcp1-mCherry or Sad1-mCherry as well as mCherry-Atb2 (α-tubulin) to visualize the SPB and spindle microtubules, respectively (Wälde and King, 2014). By relating spindle length (mCherry-Atb2 signal) to the rate of anaphase B spindle elongation (Krüger et al., 2019), we categorized anaphase B cells as early (0–5 min), mid (5–7.5 min), or late (>7.5 min).

**Figure 1.**
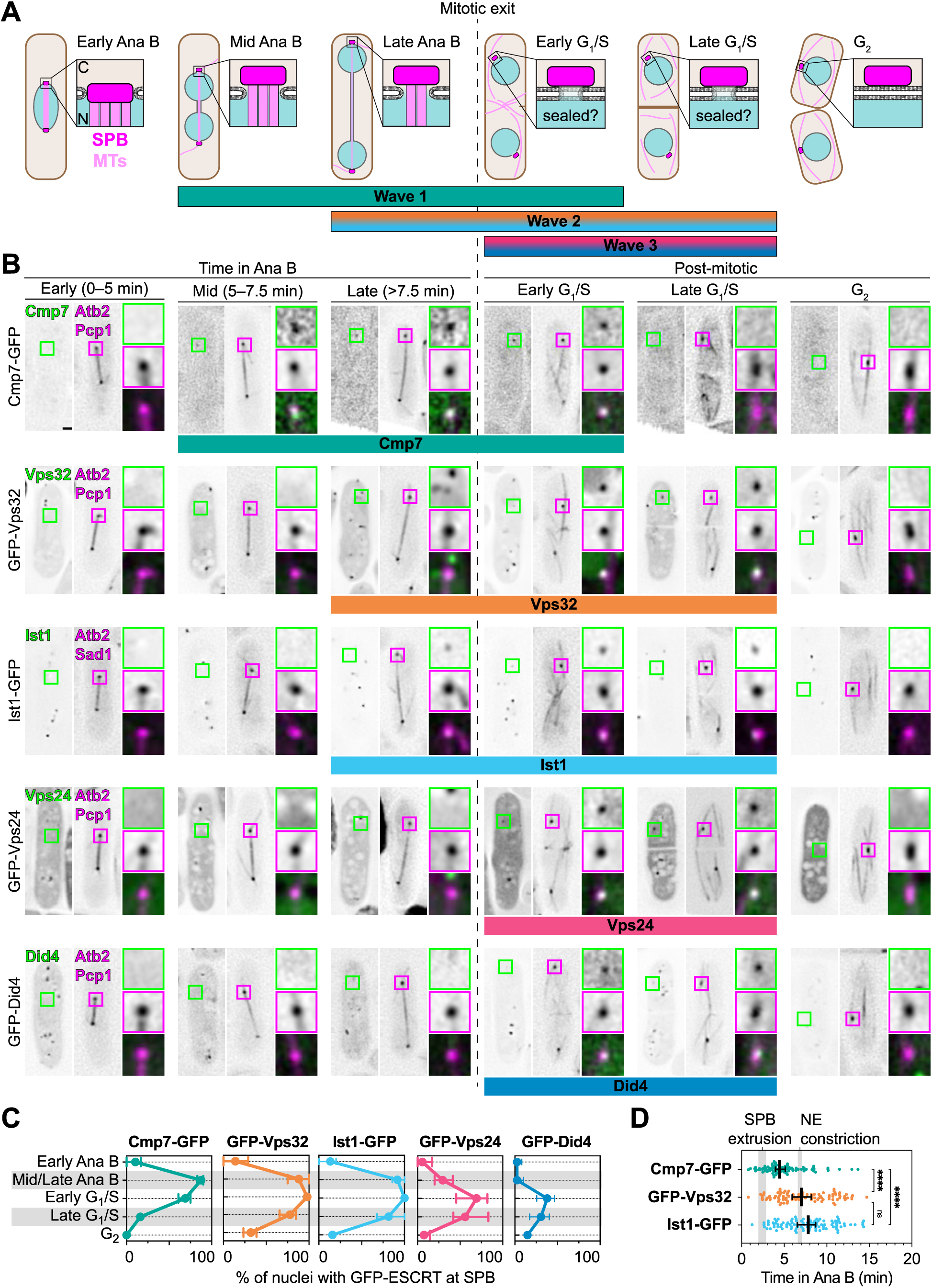
ESCRT proteins are recruited to the SPB extrusion site in three temporally distinct waves. **A:** Diagrammatic representation of *S. pombe* cell cycle from early anaphase B through G2 of the following cell cycle to establish the timeline in panel B. The spindle pole body (SPB) extrusion site from the nuclear envelope (NE) is magnified with the membrane architecture, if known, as described in Ding et al., 1997. N, nucleus (blue); C, cytoplasm (beige); SPB, dark magenta; microtubules (MTs), light magenta. Duration of the three waves of ESCRT recruitment observed in this study are indicated by labeled colored bars. Mitotic exit is indicated by the dotted vertical line. **B:** Gallery of live-cell fluorescence micrographs of GFP-tagged ESCRT proteins in *S. pombe* representative of cell populations at the indicated cell cycle stages (see key in A). In each subpanel, images of the GFP-ESCRT are on left, sum projection (over 4.8 µm) of mCherry-Atb2/Pcp1-mCherry or mCherry-Atb2/Sad1-mCherry in middle and magnifications of square region of interest (ROI) shown vertically arranged at right with (top to bottom) green, magenta and merged channels. To maximize image clarity, fluorescence intensity of magnified ROIs were independently adjusted in the linear range with the exception of the magenta channel where the gamma was adjusted to 0.05 to allow visualization of mCherry-Atb2 and Pcp1-mCherry or Sad1-mCherry without saturation. Colored bars below images indicate timing of peak ESCRT recruitment in the population (see C). Scale bar, 1 µm. Magnified ROIs, 1.6 µm wide. Cmp7-GFP, MKSP3363; GFP-Vps32, MKSP3353; Ist1-GFP, MKSP3365; GFP-Vps24, MKSP3352; GFP-Did4, MKSP3334. **C:** Plots of the percentage of nuclei with indicated GFP-ESCRT colocalized with SPB marker at indicated phases of the cell cycle. Points and error bars represent mean and range, respectively, across at least three biological replicates with at least 50 nuclei/GFP-ESCRT/replicate. Cmp7-GFP (MKSP3363), N = 408 nuclei; GFP-Vps32 (MKSP3353), N = 442 nuclei; Ist1-GFP (MKSP3365), N = 401 nuclei; GFP-Vps24 (MKSP3352), N = 699 nuclei; GFP-Did4 (MKSP3334), N = 623 nuclei. **D:** Scatter plot where each data point represents the time in anaphase B that a GFP-ESCRT was first visible colocalized with the SPB marker across three biological replicates. Bar and error bars represent mean and standard deviation of points, respectively. Vertical gray bars indicate one standard deviation of the mean across three biological replicates for Cut11-GFP (MKSP1697) focal loss (SPB extrusion, 1.96– 2.87 min) and NE constriction (6.59–7.06 min) (N = 75 cells (25 cells/replicate)). Kruskal-Wallis test performed across all datasets with Dunn’s multiple comparisons across all datasets (****, p < 0.001; n=100; at least 17 nuclei/strain/replicate).

We first determined the percentage of cells in each stage of the cell cycle where individual GFP-ESCRTs co-localized with Pcp1-mCherry or Sad1-mCherry (Figures 1B and 1C). Except for Cmp7-GFP, all GFP-ESCRTs localized to numerous cytoplasmic foci that we interpret to be endosomes (Figure 1B). Consistent with the idea that ESCRTs are only recruited to the NE when there is a NE hole, we rarely observed GFP-ESCRTs colocalized with the SPB in G_2_-phase of the cell cycle when the NE is expected to be fully sealed (Figures 1B and 1C). In contrast, except for Did2-GFP, which was never seen at the NE (Supplementary Figures S1A and S1B), all other tested ESCRT fusions colocalized with the SPB marker in cells from mid anaphase B to late G_1_/S phases (Figures 1B and 1C), which correlates with the timing of SPB extrusion from the NE (Ding *et al*., 1997). Interestingly, the peak percentage of cells when a given GFP-ESCRT accumulated at the SPB site could be temporally segregated into three waves: Cmp7 defined the first wave (Mid Ana B, “Wave 1”), Vps32 and Ist1 the second (Late Ana B, “Wave 2”,), and Vps24 and Did4 the third (G_1_/S, “Wave 3”) (Figure 1C).

To further probe the relative timing of Wave 1 and Wave 2, we performed timelapse microscopy of cells progressing through mitosis expressing Cmp7-GFP, GFP-Vps32, or Ist1-GFP. We plotted the timing of the arrival (i.e., the first frame when observed) of each GFP-ESCRT at the SPB extrusion site with respect to anaphase B progression (Figure 1D). We also placed this timeline in the context of the loss of Cut11 at the SPB, which is thought to reflect SPB extrusion, and nuclear constriction into a dumbbell shape (Figure 1D, vertical grey bars) (West et al., 1998). Using this approach, we confirmed that Cmp7-GFP was recruited to the SPB site significantly earlier than GFP-Vps32 and Ist1-GFP, which were detected with similar timing (Figure 1D). Finally, to rigorously test if we could temporally segregate the Wave 1 and 2 ESCRTs in anaphase, we examined the localization of Cmp7-GFP and Ist1-mCherry within the same cell. Indeed, we found that Cmp7-GFP appeared at the SPB extrusion site prior to recruitment of Ist1-mCherry (Supplementary Figure S1C). Thus, taken together, we conclude that there are three temporally distinct waves of ESCRT-III recruitment to the SPB extrusion site: (1) Cmp7 in mid anaphase B, (2) Vps32/Ist1 in late anaphase B, and (3) Vsp24/Did4 in the subsequent G_1_/S when Cmp7 departs (Figure 1A).

### ESCRT protein copy number at the SPB extrusion site

With perhaps one exception (Adell *et al*., 2017), there remains uncertainty regarding the range of numbers of ESCRT subunits required to execute a given membrane fission reaction. Therefore, to inform on the organization and architecture of the ESCRT polymers at the fission yeast NE, we calculated ESCRT subunit copy number at the SPB extrusion site using a semi-automated ratiometric fluorescence quantification approach. We directly related GFP-ESCRT fluorescence at the SPB site to kinetochore foci formed by a GFP-tagged kinetochore protein, Fta3, which is found at ∼37 molecules per kinetochore (for details see Methods and Supplementary Figures S2A–S2C) (Lawrimore *et al*., 2011).

Using the Fta3-GFP foci as standards, we quantified the average copy number of all ESCRT subunits recruited to the SPB extrusion site in anaphase B and G_1_/S (Table 1). Overall, there was an average of 16 ± 8 molecules of Cmp7-GFP per SPB extrusion site. The spread in the Cmp7-GFP numbers did not reflect any change in copy number during anaphase B progression or into the subsequent G_1_/S-phases (Supplementary Figures S2D and S2E). The Wave 2 ESCRT GFP-Vps32, regardless of mitotic stage, was found at a much higher copy number of 72 ± 41 molecules (Supplementary Figures S2D and S2F), suggesting an approximate 1 to 5 stoichiometry of Cmp7 to Vps32 molecules at this NE hole. The Wave 3 ESCRT subunits GFP-Vps24 and GFP-Did4 were calculated to be 77 ± 51 and 25 ± 10 molecules/focus, respectively (Supplementary Figure S2D). The one exception to an apparently stochastic variation in ESCRT subunit copy number was the Wave 2 protein Ist1-GFP, which increased in number from 25 ± 13 molecules in anaphase B to 34 ± 18 into G_1_/S (Supplementary Figure S2D). Consistent with an increase in Ist1 copy number through anaphase B, there was a positive correlation between Ist1-GFP copy number at individual NE holes with anaphase B progression (Supplementary Figure S2G). This incremental accumulation of an ESCRT was not observed for other ESCRTs in our work or that of others at ILVs (Adell *et al*., 2017), suggesting that it could be a behavior unique to Ist1.

**Table 1.** Average copy number for ESCRT proteins at the NE. Copy number is shown for all examined ESCRTs by cell cycle. Copy number at intralumenal vesicle (ILV) formation at endosomes as reported in Adell et al. (2017). *n.r.,* not recruited. *n.m.,* not measured.

| Protein | Mitotic NE hole |  |  |  |  |  | ILV<br>Formation<sup>1</sup> |
| --- | --- | --- | --- | --- | --- | --- | --- |
|  | Ana B |  | G<sub>1</sub>/S |  | All |  |  |
|  | Mean ± SD | N | Mean ± SD | N | Mean ± SD | N | Mean |
| Cmp7 | 16 ± 8 | 88 | 15 ± 8 | 108 | 15 ± 8 | 196 | *n.r.* |
| Vps32 | 75 ± 45 | 33 | 69 ± 37 | 35 | 72 ± 41 | 68 | 75–200 |
| Ist1 | 25 ± 13 | 67 | 34 ± 18 | 80 | 30 ± 17 | 147 | *n.m.* |
| Vps24 | *n.m.* |  | 77 ± 51 | 24 | 77 ± 51 | 24 | 15–50 |
| Did4 | *n.r.* |  | 25 ± 10 | 26 | 25 ± 10 | 26 | *n.m* |

### Cmp7 is required for ESCRT recruitment to the NE

The sequential timing of ESCRT recruitment waves to the NE suggests a dependency of the later wave ESCRTs on those preceding them. To test this idea, we next determined the co-dependency of ESCRT protein recruitment to the mitotic NE hole. As predicted from prior work in budding yeast (Webster *et al*., 2016) and cultured human cells (Gu *et al*., 2017), the recruitment of Cmp7-GFP to the SPB site required Heh1/Lem2 but not its paralog Heh2/Man1 (Figures 2A and 2B). Consistent with the observed sequential nature of ESCRT recruitment, Wave 2 or 3 ESCRTs failed to be recruited to the NE in the absence of the Wave 1 Cmp7 (Figure 2C). Thus, as has been observed in other contexts (Gu *et al*., 2017; Olmos *et al*., 2016; Thaller *et al*., 2019; Vietri *et al*., 2015), Cmp7 is required for all ESCRT-III recruitment to the NE.

**Figure 2.**
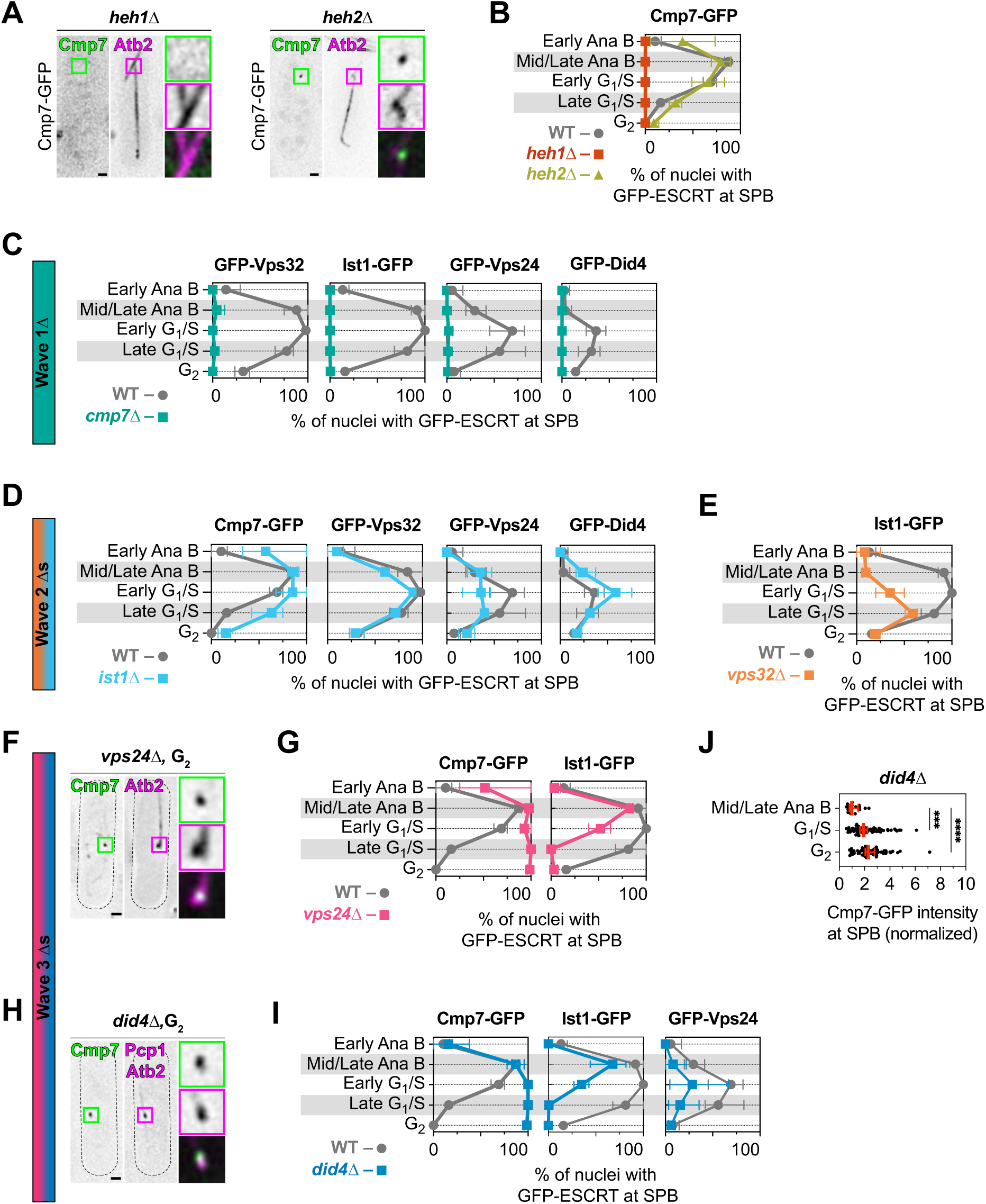
Cmp7 is required for ESCRT-III recruitment to the NE and is replaced by Wave 3 ESCRTs. **A:** Fluorescence micrographs of Cmp7-GFP and mCherry-Atb2 in representative *heh1Δ* (left) and *heh2Δ* (right) cells in anaphase B. In each subpanel, images of the Cmp7-GFP are on the left, sum projection (over 4.8 µm) of mCherry-Atb2 in middle and magnifications of square ROI shown vertically arranged at right with (top to bottom) green, magenta and merged channels. Fluorescence intensity of magnified ROIs were independently adjusted in the linear range. Scale bar, 1 µm. Magnified ROIs, 1.6 µm wide. **B:** Plots of the percentage of nuclei with Cmp7-GFP foci colocalized with the SPB site in WT (gray circles), *heh1Δ* (red squares), and *heh2Δ* (yellow triangles) strains in indicated phases of the cell cycle. Points and error bars represent mean and range, respectively, across at least three biological replicates with at least 50 nuclei/Cmp7-GFP/replicate. Cmp7-GFP, WT (data from Figure 1C), Cmp7-GFP, *heh1Δ* (MKSP3259), N = 203; Cmp7-GFP, *heh2Δ* (MKSP3275), N = 275 nuclei. **C:** As in B but with the indicated GFP-ESCRTs in WT (gray circles) or *cmp7Δ* (green squares) cells. Points and error bars represent mean and range, respectively, across at least three biological replicates with at least 50 nuclei/GFP-ESCRT/replicate. All WT data from Figure 1C; GFP-Vps32, *cmp7Δ* (MKSP3354), N = 628 nuclei; Ist1-GFP, *cmp7Δ* (MKSP3447), N = 384 nuclei; GFP-Vps24, *cmp7Δ* (MKSP3355), N = 702 nuclei; GFP-Did4, *cmp7Δ* (MKSP3366), N = 372 nuclei. **D:** As in B but with the indicated GFP-ESCRTs in WT (gray circles) or *ist1Δ* (blue squares) cells. Points and error bars represent mean and range, respectively, across at least three biological replicates with at least 50 nuclei/GFP-ESCRT/replicate. All WT data from Figure 1C; Cmp7-GFP, *ist1Δ* (MKSP3429), N = 418 nuclei; GFP-Vps32, *ist1Δ* (MKSP3449), N = 550 nuclei; GFP-Vps24, *ist1Δ* (MKSP3669), N = 323 nuclei; GFP-Did4, *ist1Δ* (MKSP3668), N = 455 nuclei. **E:** As in B but with Ist1-GFP in WT (gray circles) or *vps32Δ* (orange squares) cells. Ist1-GFP, WT (data from Figure 1C), Ist1-GFP, *vps32Δ* (MKSP3504), N = 382 nuclei. **F:** As in A but in *vps24Δ*, G2 cells. **G:** As in B but with the indicated GFP-ESCRTs in WT (gray circles) or *vps24Δ* (red squares) cells. All WT data from Figure 1C; Cmp7-GFP, *vps24Δ* (MKSP3387), N = 236 nuclei; Ist1-GFP, *vps24Δ* (MKSP3544), N = 378 nuclei. **H:** As in A but in *did4Δ*, G2 cells. Gamma was adjusted to 0.05 to allow visualization of mCherry-Atb2 and Pcp1-mCherry without saturation. **I:** As in B but with the indicated GFP-ESCRTs in WT (gray circles) or *did4Δ* (blue squares) cells. All WT data from Figure 1C; Cmp7-GFP, *did4Δ* (MKSP3381), N = 323 nuclei; Ist1-GFP, *did4Δ* (MKSP3496), N = 376 nuclei; GFP-Vps24, *did4Δ* (MKSP3511), N = 438 nuclei. **J:** Plot of the fluorescence intensity of Cmp7-GFP at the SPB site in *did4Δ* cells (MKSP3381) at indicated cell cycle stages normalized to median mid/late anaphase B value for each biological replicate. Bar and error bars represent mean and standard deviation of points, respectively. Kruskal-Wallis test performed across all datasets with Dunn’s multiple comparisons across all dataset (****, p < 0.001; N=160, at least 45 cells/replicate).

We next tested the requirement of Wave 2 ESCRTs for recruitment of other GFP-ESCRTs. As shown in Figure 2D, Ist1 is entirely dispensable for all GFP-ESCRT recruitment to the SPB extrusion site. Interestingly, Ist1-GFP recruitment is compromised but not completely abrogated in the absence of its Wave 2 partner, Vps32, appearing both later and in a smaller percentage of cells compared to WT (Figure 2E). Thus, although Cmp7 is essential for any ESCRT-III subunit recruitment, there may not be a strict assembly order for other downstream filament assembly events.

### Cmp7 is replaced by Wave 3 ESCRT assemblies in G_1_/S

As our temporal analysis suggests progressive remodeling of the ESCRT complement at the NE hole in concert with the cell cycle, we next tested whether the inability to recruit the final wave of ESCRTs impacted the dynamics of the upstream Wave 1 or 2 ESCRTs. Strikingly, even though the Wave 3 ESCRTs Vps24 and Did4 were found at the SPB site when the majority of Cmp7 had departed in most WT cells (Figure 1), we found that Cmp7-GFP recruitment was dysregulated in *vps24Δ* and *did4Δ* cells, remaining at the NE throughout G_1_/S and even G_2_ (Figures 2F–2I). Moreover, Cmp7-GFP over-accumulated at the NE in *vps24Δ* and *did4Δ* cells with a median 2-fold greater fluorescence than in WT cells (Figure 2J). Interestingly, although the initial recruitment of the Wave 2 ESCRT Ist1-GFP was normal in *vps24Δ* and *did4Δ* cells, Ist1-GFP failed to persist with the same kinetics observed in WT cells (Figures 2G and 2I). Moreover, although still observed in a small percentage of SPB extrusion sites, Vps24 recruitment was largely dependent on its Wave 3 partner, Did4 (Figure 2I). Thus, we interpret these data in a model in which the inability to complete ESCRT remodeling during G_1_/S drives a reversal of the ESCRT order into an aberrant Wave 1 state with persistent Cmp7-GFP recruitment.

### ESCRT waves correlate to changes in NE hole morphology

To help understand ESCRT function from anaphase to G_1_/S, we correlated the ESCRT recruitment timeline to the ultrastructure of the NE hole at the SPB extrusion site using correlative light and electron microscopy (CLEM) and electron tomography (Figure 3). We used Pcp1-mCherry and Cdc7-GFP, the latter of which transiently localizes to mitotic/G_1_ spindle poles (Sohrmann et al., 1998), to help inform the location of the SPB extrusion site and the cell cycle stage of embedded cells. At the correlated location of these SPB markers, we identified the SPB plaque as an electron-dense region in the cytoplasm apposing the nucleus (Figures 3A–3D, red outline). We also observed an electron-lucent region of cytosol surrounding the SPB plaques that excluded ribosomes (Figures 3A–D, orange outline).

**Figure 3.**
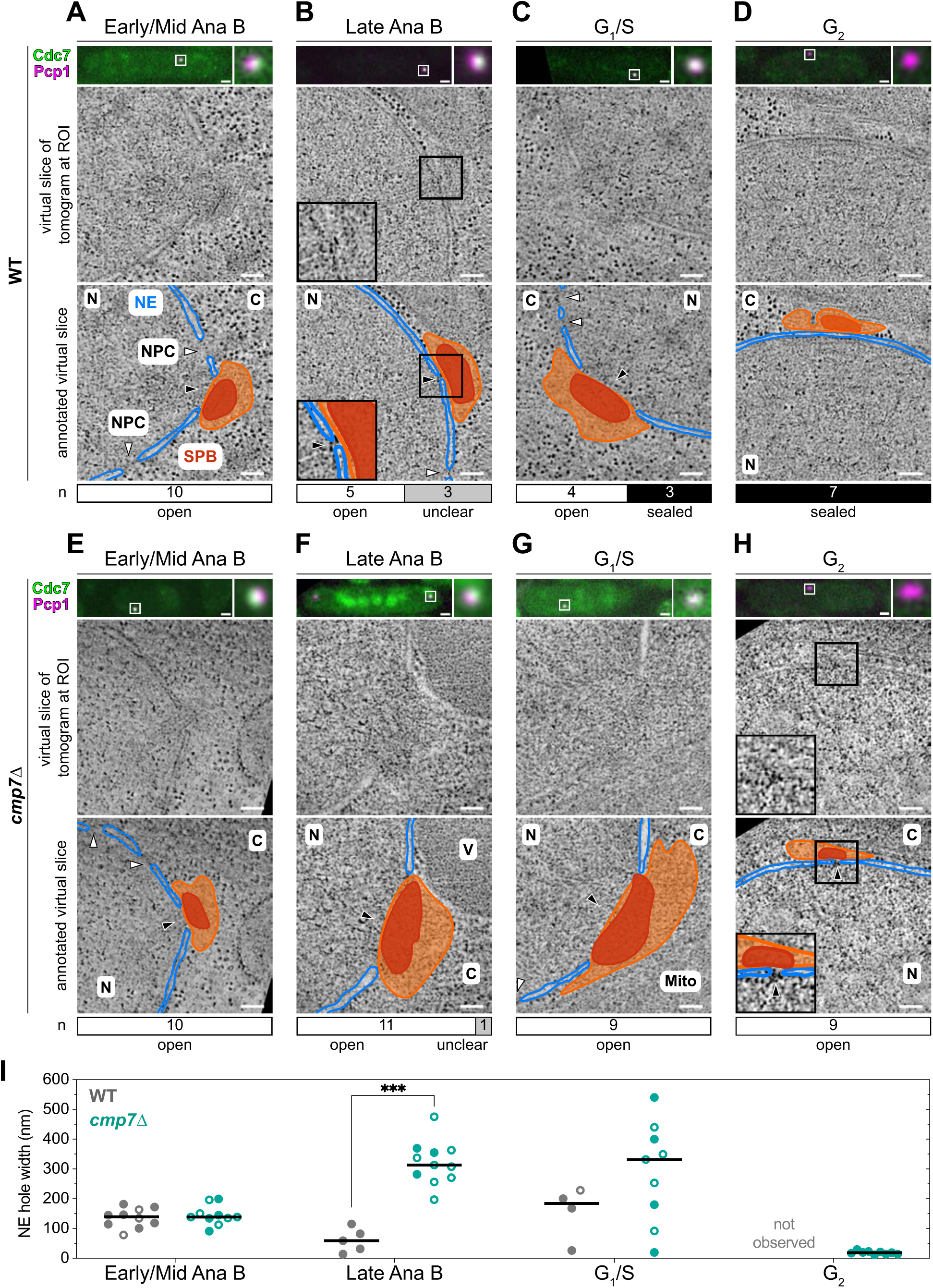
Cmp7 is required to restrict the size of the NE hole in anaphase and seal the NE in G1/S. **A–D:** Top, fluorescence micrograph of a thick section of a resin-embedded yeast strain (MKSP3180) expressing Cdc7-GFP (green) and Pcp1-mCherry (magenta) prepared for correlative light and electron microscopy (CLEM) and electron tomography (ET). The indicated cell cycle stages were determined by a combination of fluorescence marker presence/localization and ultrastructural features. The white square in the fluorescence micrographs indicates regions where the electron tomogram was collected and is shown magnified at the right. Middle and bottom panels are a virtual slice taken from the electron tomogram shown without (middle) and with (bottom) manual annotation of features. NE, blue, SPB plaques, red, electron-lucent ribosome exclusion zone extending beyond SPB plaques, orange. White arrowheads, NPC. Black arrowheads, SPB extrusion hole. N, nucleus; C, cytoplasm. Scale bars: fluorescence micrographs, 1 µm; virtual slices from electron tomogram, 100 nm. **E–H:** CLEM and ET for *cmp7Δ* cells (MKSP3243), prepared and displayed as in A. Mito, mitochondria; V, vacuole. **I:** Scatter plot of NE hole widths in tomograms of WT (gray circles) and *cmp7Δ* (green circles). Solid circles represent exactly determined diameters while open circles represent minimum values in cases where the full diameter was not visualized in the volume. Black bars represent median. Mann-Whitney test performed between WT and *cmp7Δ* for each cell cycle phase individually (***, p < 0.0001; all others, p ≥ 0.05). WT cells, MKSP3180, N = 14 nuclei; *cmp7Δ* cells, MKSP3243, N = 39 nuclei.

The onset of Wave 1 ESCRT recruitment occurs in mid anaphase B; in early/mid anaphase B cells examined (n=10) we observe a median value of the fenestration the NE apposing the SPB of 140 nm (Figure 3A, black arrowhead; Figure 3I); spindle microtubules passed through this hole in select virtual slices of the entire tomogram (Movie 1). Interestingly, we observed a narrowing of the NE hole to a median of 59 nm in late anaphase B cells, alongside a reduction of microtubule number (Figure 3B, black arrowhead; Figure 3I, n=5) (Movie 2) (Ding *et al*., 1997). These data suggest that the recruitment of the Wave 2 ESCRTs temporally correlates with a constriction of the NE hole from ∼140 to ∼60 nm. Subsequently, in G_1_/S cells there was either a relaxation of the NE hole to a median of 184 nm (n=4; Figure 3C, black arrowhead and 3I) or cells that had completed NE sealing (n=3), suggesting that the timing of Wave 3 corresponds to the act of NE sealing. Consistent with this interpretation, the NE membranes were continuous in all 7 G_2_-phase cells examined (Figures 3D and 3I). Thus, each wave of ESCRT recruitment corresponds to distinct morphological changes of the NE hole; Wave 2 corresponds to a transient NE constriction while Wave 3 represents a timepoint that is suggestive of a role in NE sealing.

### ESCRTs restrict the size of the NE hole during mitosis

To directly evaluate whether ESCRTs impact the observed NE hole dynamics, we performed CLEM on *cmp7Δ* cells, which fail to recruit all ESCRT-III proteins (Figure 2C). Loss of *CMP7* did not impact the morphology of the NE fenestration in early/mid anaphase B cells, which were similar to those in WT cells (n=10; Figures 3E and 3I; Movie 3). In striking contrast, late anaphase B *cmp7Δ* cells (n=12) exhibited massive NE gaps with a median width of 313 nm and a high variation (range of 197–475 nm; Figures 3F and 3I; Movie 4), an approximate five-fold increase compared to WT cells at the same stage. Thus, we conclude that Cmp7 is required to restrict the diameter of the mitotic NE hole during anaphase B. Given that this state temporally precedes the recruitment of Wave 3 ESCRTs (Figure 1), this function is likely carried out by Wave 1 and Wave 2 ESCRTs.

### Cmp7 is required but must be remodeled to Wave 3 ESCRTs to promote NE sealing

Interestingly, in G_1_/S when some WT NE holes had been sealed, NE gaps that ranged from 20 to 540 nm (median, 332 nm) were observed in all imaged *cmp7Δ* cells; n=6, Figures 3G and 3I). Although not statistically different, the large spread of values in individual *cmp7Δ* cells suggests that the aberrantly large fenestrations arising in anaphase B persist into the subsequent G_1_/S. Most strikingly, G_2_ *cmp7Δ* cells (n=9, Figure 3H) retained a discernable but small median 19 nm NE fenestration (Figure 3I), suggesting that an ESCRT-independent mechanism ultimately constricted, but failed to seal, the NE.

As our light microscopy data indicated that Cmp7 aberrantly persists in cells lacking Wave 3 ESCRTs (Figures 2F–2J), we also interrogated the morphological consequences of reversion to an aberrant Wave 1 assembly state by applying CLEM and electron tomography to G_2_ *did4Δ* cells (Figures 4A and 4B). At the majority (8/13) of Cmp7-GFP NE foci we selected, we correlated the fluorescence to a median 21 nm hole in the NE (Figure 4C), suggesting that the persistent Wave 1 state also disrupts NE sealing; in the other 5 samples, we could not unambiguously observe a NE hole (Figures 4A and 4B). Unlike in *cmp7Δ* cells, however, in *did4Δ* cells we found that the persistent NE hole was filled with densely stained material (Figure 4A), which we interpret to be Cmp7-GFP, although other, yet to be defined factors may also be present. Interestingly, although Cmp7-GFP persisted at the SPB extrusion site throughout G_2_ in *did4Δ* cells, it disappeared with the onset of the subsequent SPB insertion cycle (Figure 4D). These data suggest a mechanism to “reset” the SPB site for another round of SPB insertion, extrusion and NE sealing that might entail clearing aberrant ESCRT assemblies from the NE. Thus, we conclude that Cmp7’s role in initiating ESCRT-III assembly is required to close the constricted, 20 nm NE fenestration in G_2_. Moreover, the loss of Wave 3 ESCRTs leads to the accumulation of Cmp7 in NE holes that fail to seal, highlighting that it is the replacement of the initial Wave 1/Wave 2 ESCRT assemblies to the Wave 3 ESCRT state that ultimately drives the sealing reaction.

**Figure 4.**
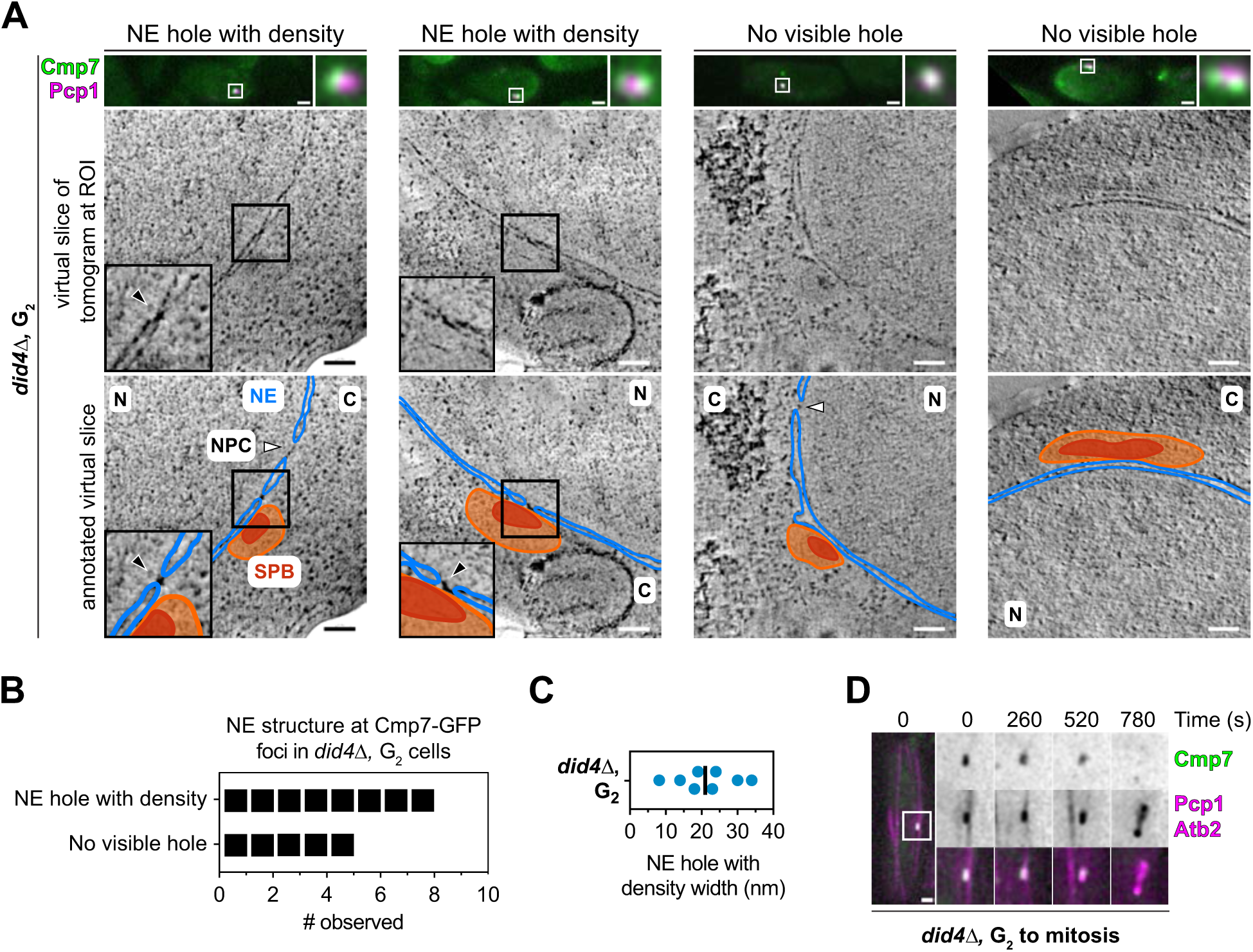
Did4 is required to remove Cmp7 to seal the NE. **A:** Gallery of four representative electron tomograms showing either NE holes filled with electron density (left) or no visible NE holes (right). In each sub panel: Top, fluorescence micrograph of a thick section of a resin-embedded *did4Δ* strain (MKSP3381) in G2 expressing Cmp7-GFP (green) and Pcp1-mCherry (magenta) prepared for CLEM and ET. The white square in the fluorescence micrographs indicates the region where the electron tomogram was collected and is shown magnified at right. Middle and bottom panels are a virtual slice taken from the electron tomogram shown without (middle) and with (bottom) manual annotation of features. NE, blue, SPB plaques, red, electron-lucent ribosome exclusion zone extending beyond SPB plaques, orange. White arrowheads, NPC. Black arrowheads, NE hole with density. N, nucleus; C, cytoplasm. Scale bars: fluorescence micrographs, 1 µm; virtual slices from electron tomogram, 100 nm. **B:** Plot of the number of occurrences (black squares) of the observed NE phenotypes from A. MKSP3381, N = 13 nuclei. **C:** Scatter plot of NE hole widths (nm) observed in *did4Δ*, G2 cell tomograms as in A. Black bar represents median. MKSP3381, N = 8 nuclei. **D:** Fluorescence micrographs of a timelapse series (top, seconds representing the transition from G2 into mitosis) of a *did4Δ* cell expressing Cmp7-GFP (green), Pcp1-mCherry (magenta), and mCherry-Atb2 (magenta). The white square in the overview micrograph indicates the ROI magnified in the image series vertically arranged at right with (top to bottom) green, magenta and merged channels. Representative cell (MKSP3381) from three biological replicates of imaging N = 180 cells, at least 60 cells/replicate.

### Cmp7 is dispensable for maintaining nucleocytoplasmic compartmentalization

The persistence of NE holes at the SPB extrusion site throughout the cell cycle in *cmp7Δ* cells, and in particular the massive NE fenestrations observed during mitosis, prompted an evaluation of nucleocytoplasmic compartmentalization in these conditions. As the steady-state distribution of NLS-containing reporters, and by inference NPC function, are not affected by the loss of *CMP7* (Gu *et al*., 2017), we investigated the passive nuclear exclusion of large reporters lacking nuclear import or export signals. Specifically, we tested the distribution of GFP fused to either three (MGM2) or five (MGM4) maltose-binding proteins; while MGM2 is able to diffuse across NPCs, the larger MGM4 construct is effectively excluded from the nucleus (Figure 5A and Supplementary Figure S3A) (Popken et al., 2015). Consistent with this, most MGM2 cells had nuclear to cytoplasmic ratios (N/C ratios) >0.75 and most MGM4 cells had N/C ratios <0.75; we therefore used N/C ratio = 0.75 as a cut-off to gauge whether cells expressing MGM4 displayed a defect in their nucleocytoplasmic compartmentalization (Supplementary Figure S3A). Using this framework, we found that *cmp7Δ* cells displayed no detectable loss of nuclear compartmentalization (Figure 5A).

**Figure 5.**
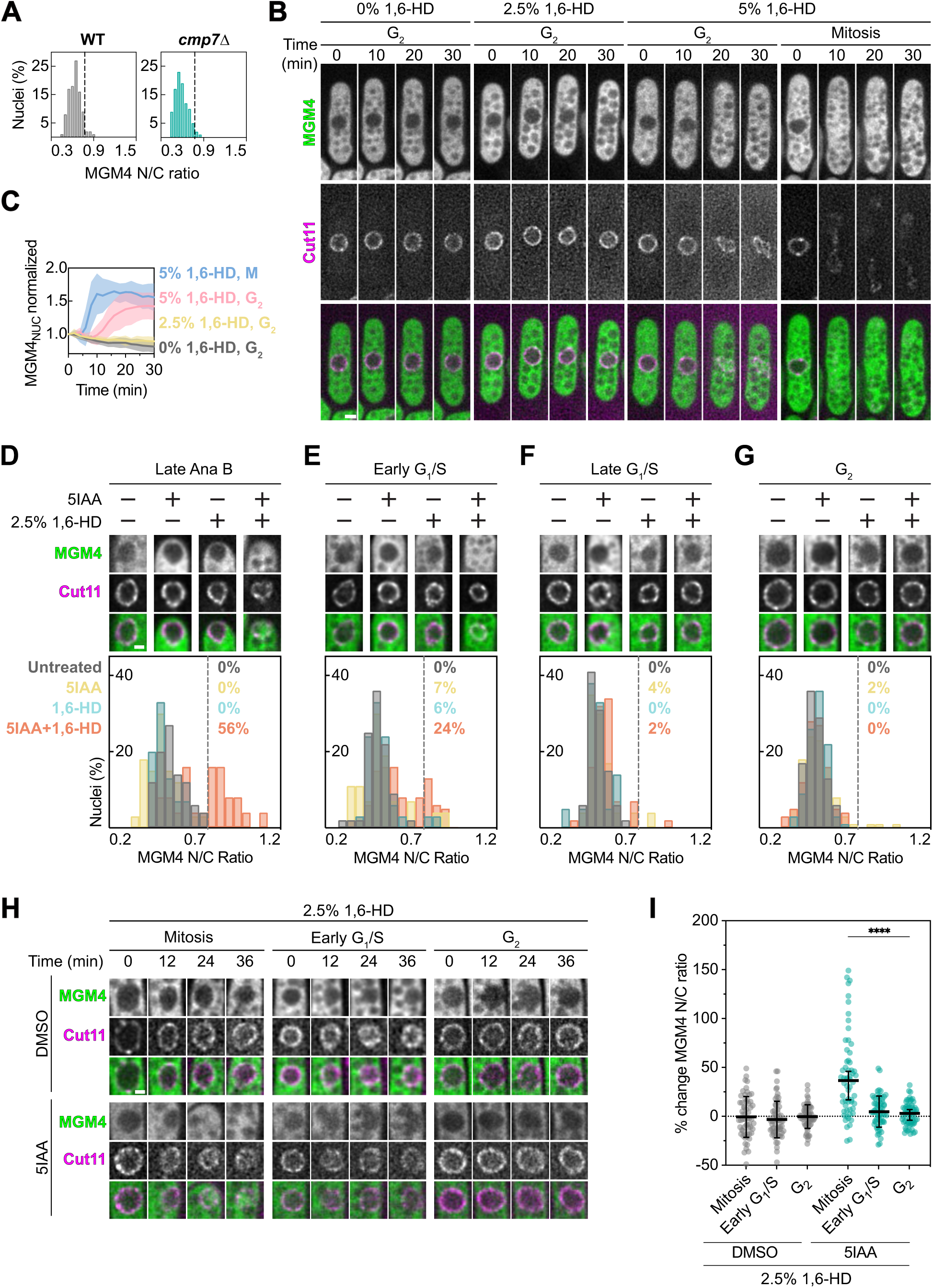
Cmp7 and a 1,6 hexanediol-sensitive diffusion barrier ensure nucleocytoplasmic compartmentalization during mitosis. **A:** Histogram relating nucleoplasmic and cytoplasmic GFP fluorescence of MGM4 in a N/C ratio of individual WT and *cmp7Δ* cells across three biological replicates. Size between bins, 0.05. Dotted line at 0.75 bin. WT (MKSP3345 and MKSP3488), N = 240 nuclei; *cmp7Δ* (MKSP3535), N= 177 nuclei. **B:** A time-lapse series of deconvolved fluorescence micrographs of cells (MKSP3699) expressing MGM4 and Cut11-mCherry. MGM4 (green, top), Cut11-mCherry (magenta, middle) and merged image (bottom) shown. Cells were treated with 0%, 2.5% or 5% 1,6-hexanediol (1,6-HD) as indicated. For G2 cells, time 0 indicates the start of imaging after the delivery of 1,6-HD by microfluidics. For mitotic cells, only cells that went through mitosis after 1,6-HD treatment were considered. Across the population, the constriction of the NE into a dumbbell shape (∼10 min) was used to align individual traces in C. Scale bar, 1 µm. Representative of 20 cells per condition with at least 5 cells per condition for each of three biological replicates. **C:** Plot of the nuclear (NUC) fluorescence of MGM4 through timelapse imaging experiments as shown in B normalized to the initial timepoint for each individual nucleus of a cell (MKSP3699, untreated). Solid lines represent average of 20 cells per condition with at least 5 cells per condition for each of three biological replicates, with the shaded region representing the standard deviation. M = mitosis. **D–G:** Asynchronous cells (MKSP3699) expressing MGM4, Cut11-mCherry, Cmp7-AID and OsTIR1F74A were treated for 1 h with DMSO or 5-adamantyl-IAA (5IAA) followed by EMM5S media with or without 2.5% 1,6-HD for 45 to 90 min—treatments indicated by pluses—before imaging. Post-imaging, cells were categorized by cell cycle stage and analyzed. At the top of each panel are representative images of an ROI that encompasses a nucleus with MGM4, Cut11-mCherry and merged images arranged vertically. Scale bar, 1 µm. At bottom, histograms of N/C ratios of MGM4 fluorescence of individual cells in untreated (gray), 5IAA (yellow), 2.5% 1,6-HD (teal), and both 5IAA and 2.5% 1,6-HD (orange). Percent of values above 0.75 bin (dotted gray line) for each reporter indicated. Size between bins, 0.05. Quantification was performed for three biological replicates N = 955, with at least 197 cells/condition, and at least 50 cells/biological replicate. **H:** DMSO or 5IAA-treated cells (MKSP3699) at indicated cell cycle stages expressing MGM4, Cut11-mCherry, Cmp7-AID and OsTIR1F74A were subjected to timelapse imaging after the addition of 2.5% 1,6 HD delivered by microfluidics. For G2 and G1/S cells, time 0 indicates the start of imaging after the delivery of 1,6-HD by microfluidics. For mitotic cells, only cells that went through mitosis after 1,6-HD treatment were considered. Images of representative nuclei at each timepoint are shown arranged vertically with MGM4 (green) at top, Cut11-mCherry (magenta) in middle and merge at bottom. Scale bar, 1 µm. Representative of N=360, 20 cells/replicate/cell cycle phase/treatment. **I:** Scatterplot of the percent change of MGM4 N/C ratios of individual cells (MKSP3699, either DMSO or 5IAA-treated) expressing MGM4, Cut11-mCherry, Cmp7-AID and OsTIR1F74A from start to end of timelapse imaging after addition of 2.5% 1-6 HD. Bars and error bars represent mean and standard deviation, respectively. Kruskal-Wallis test performed across all cell cycle phases with Dunn’s multiple comparisons for each data set against G2 for the corresponding treatment (****, p < 0.0001; ns, p ≥ 0.05). N=360, 20 cells/replicate/cell cycle phase/treatment.

### Cmp7 cooperates with a diffusion barrier to maintain the nuclear compartment

The observation that deletion of *CMP7* did not appreciably impact nucleocytoplasmic compartmentalization despite the presence of large NE fenestrations was, on the surface, surprising. It should be considered, however, that even in WT cells there is a period (∼14 minutes) after SPB extrusion when the nuclear compartment appears intact despite the presence of a NE fenestration of at least ∼60 nm but up to 228 nm beneath the SPB (Figure 3). Thus, there must be additional, unidentified diffusion barrier(s) (beyond the NPC) capable of maintaining nucleocytoplasmic compartmentalization in the face of a persistent NE hole. To begin to test this idea, we employed 1,6-hexanediol (1,6-HD), an aliphatic alcohol that disrupts weak hydrophobic protein-protein interactions including that of FG-repeat containing nucleoporins that contribute to the NPC diffusion barrier (Ribbeck and Görlich, 2002; Riquelme Barrientos et al., 2023; Shulga and Goldfarb, 2003). We observed that 5%, but not 2.5%, 1,6-HD treatment led to the equilibration of MGM4 fluorescence across the NE in about 20 minutes in WT G_2_ cells (Figures 5B and 5C), indicating disruption of the NPC diffusion barrier. Interestingly, upon 5% 1,6-HD addition MGM4 fluorescence equilibrated approximately twice as fast in mitotic cells (Figures 5B and 5C). This observation hints that either the NPC diffusion barrier is weakened during mitosis or there is an additional proteinaceous NE diffusion barrier required during mitosis that is also sensitive to 5% 1,6-HD.

Building on a model where there is a mitotic-specific NE diffusion barrier, we reasoned that such a barrier would be vulnerable to the loss of Cmp7, which leads to a much larger NE hole in anaphase B (Figures 3F and 3I). To test this, we turned to an auxin-induced degron (AID) to quickly deplete an endogenously expressed Cmp7-AID fusion, which allowed us to specifically test the impact of the loss of Cmp7 during mitosis (Kanke et al., 2011; Zhang et al., 2022). Cmp7-AID was efficiently degraded within 15 minutes of treatment with 5-adamantyl-IAA (5IAA) (Supplementary Figure S3B). Importantly, depletion of Cmp7-AID recapitulates phenotypes observed in *cmp7Δ* cells, for example abolishing recruitment of Ist1-GFP to the NE (Supplementary Figure S3C). Further, like *cmp7Δ*, there was no discernable change in MGM4 N/C fluorescence ratios in 5IAA-treated Cmp7-AID cells (Figures 5D–5G).

We therefore sought to specifically disrupt the putative mitotic diffusion barrier in Cmp7-AID depleted cells with 2.5% 1,6-HD, which is insufficient to disrupt the NPC diffusion barrier even during mitosis (Figures 5B–5D, and 5H). Strikingly, treatment of Cmp7-AID depleted cells with 2.5% 1,6-HD resulted in nuclear entry of MGM4 in 56% of late anaphase B cells and 24% of early G_1_/S cells but did not impact MGM4 distribution in G_2_ cells (Figures 5D–5G). To confirm that the nuclear entry of MGM4 occurred specifically during mitosis under conditions of Cmp7 depletion, we monitored MGM4 localization in Cmp7-AID cells (with or without 5IAA) alongside the temporally controlled delivery of 2.5% 1,6-HD in a microfluidic chamber (Figures 5H and 5I). As shown in Figure 5H, only mitotic cells depleted of Cmp7-AID in the presence of 2.5% 1,6-HD allowed entry of MGM4 into the nucleus, which we quantified as the percent change in N/C fluorescence ratios of individual cells from the beginning to the end of the experiment (Figure 5I). Together, these data suggest that Cmp7 contributes to a mitotic diffusion barrier independent of NPCs that is distinct from its role in NE membrane sealing.

### *CMP7* genetically interacts with genes encoding SPB components

We next sought to identify the protein(s) that help form a diffusion barrier capable of preventing macromolecules from transiting a NE hole as large as ∼500 nm. Based on the CLEM data, which showed the SPB and a zone of ribosome exclusion covering even the largest NE holes in *cmp7Δ* cells (Figure 3), SPB components were the most likely candidates. We conducted a genetic interaction screen between genes that encode proteins that are either constitutively or transiently (during mitosis) associated with the SPB and *CMP7* (Jaspersen, 2021; Wälde and King, 2014) (Figure 6A, B and Supplementary Figure S4). Of the 17 genetic backgrounds with perturbations to the SPB, 12 were synthetically sick when combined with the *cmp7ϕ..* allele and one was synthetically lethal. The remaining four alleles that did not rely on *CMP7* for fitness affected proteins that are peripherally associated with the SPB rather than residing in the core structure. Importantly, we find genetic interactions not just between *CMP7* and genes encoding constitutive SPB proteins, but with genes that encode proteins that transiently associate with the SPB to ensure proper mitotic progression (*BRR6*, *CUT11*, and *PLO1*) (Tamm et al., 2011; Wälde and King, 2014; West *et al*., 1998). Thus, growth (and in some cases viability) of cells with a compromised SPB is dependent on *CMP7*.

**Figure 6.**
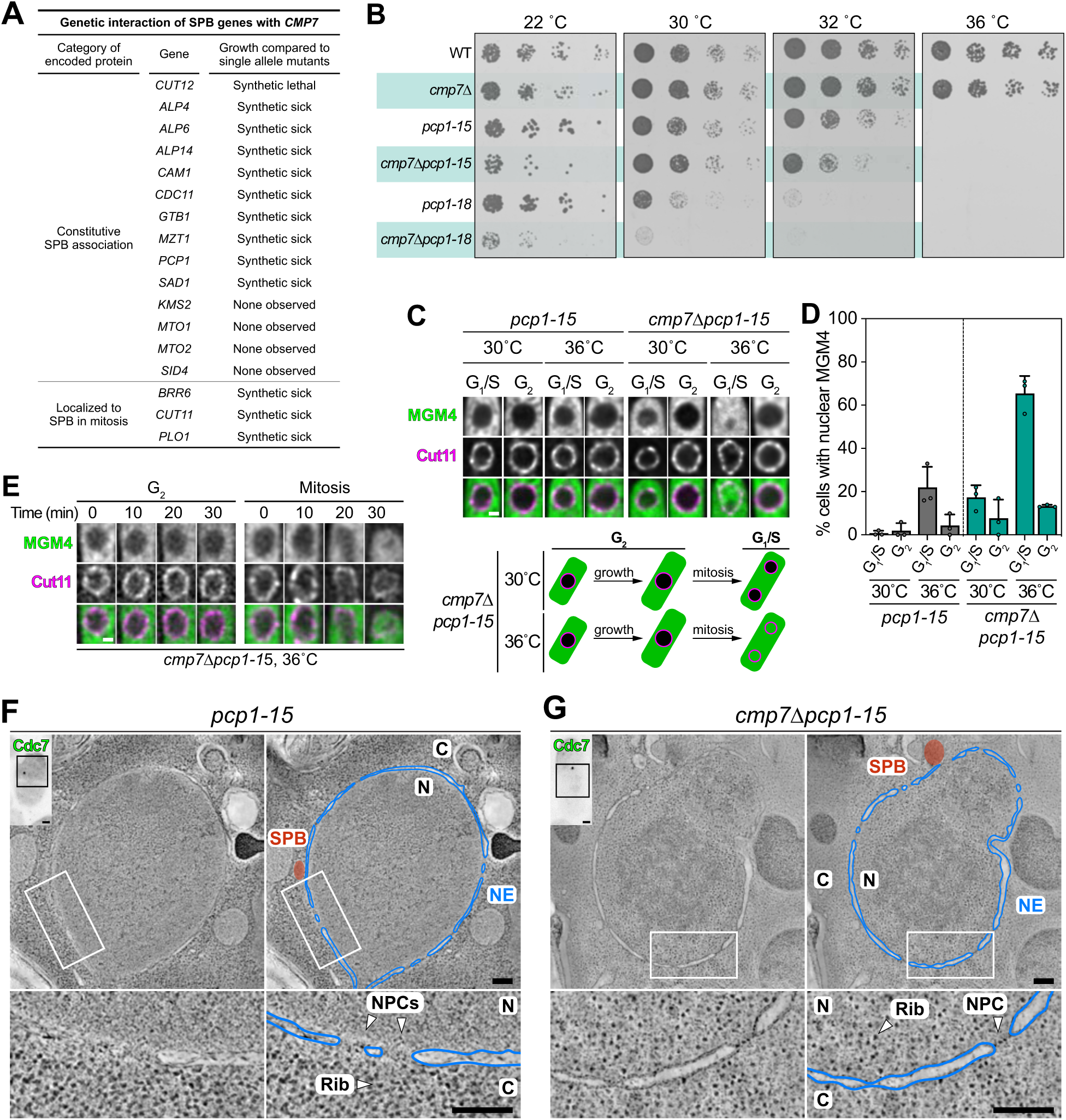
*CMP7* is required for robust fitness and nucleocytoplasmic compartmentalization in the context of perturbed SPB protein function. **A:** Summary table of observed synthetic genetic interactions from plate growth assay data shown in Supplementary Figure S4. **B:** *CMP7* is required for fitness of strains with *PCP1* mutant alleles. The indicated strains were serially diluted 10-fold and plated on YE5S plates incubated at the indicated temperatures. Plates were imaged after 2 days (30, 32, and 36°C) or 5 days (22°C). WT, MKSP399; *cmp7Δ*, MKSP3340; *pcp1-15*, MKSP280; *cmp7Δpcp1-15*, MKSP3420; *pcp1-18*, MKSP281; *cmp7Δpcp1-18*, MKSP3425. **C:** Fluorescence micrographs of representative nuclei are shown arranged vertically with MGM4 (green) at top, Cut11-mCherry (magenta) in middle and merge at bottom of either *pcp1-15* (MKSP3812) or *cmp7Δpcp1-15* (MKSP3811) strains. Cells were grown at 30°C then imaged directly or following 3 h at 36°C. Scale bar, 1 µm. Cartoon model of loss of nuclear integrity as it occurs mostly in post-mitotic G1/S *cmp7Δpcp1-15* cells grown at 36°C for 3 h. **D:** Plot of the percentage of *pcp1-15* (gray bars, MKSP3812) or *cmp7Δpcp1-15* (teal bars, MKSP3811) MGM4-expressing cells with N/C fluorescence ratios greater than 0.75 reflecting a loss of nucleocytoplasmic compartmentalization. Points are the mean from each biological replicate, while bars and error bars represent total mean and standard deviation, respectively. Points represent average of at least 30 nuclei/cell cycle phase/strain/replicate, N = 977. **E:** Fluorescence micrographs of representative nuclei are shown arranged vertically with MGM4 (green) at top, Cut11-mCherry (magenta) in middle and merge at bottom from individual *cmp7Δpcp1-15* cells (MKSP3811) in G2 or through a timecourse (in minutes) after a temperature shift from 30°C to 36°C. Scale bar, 1 µm. **F, G:** Top-left inset, fluorescence micrograph of a thick section of a resin-embedded yeast strain (MKSP4107, F; MKSP4106, G) expressing Cdc7-GFP (green) prepared for CLEM and ET following 3 h of incubation at 36°C. The black square in the fluorescence micrographs indicates regions where the electron tomogram was collected. Left and right panels are a virtual slice taken from the electron tomogram shown without (left) and with (right) manual annotation of features. White square shows magnified region shown in bottom panels. NE, blue, SPB region, red. White arrowheads, NPC. N, nucleus; C, cytoplasm; Rib, ribosomes. Scale bars: fluorescence micrographs, 1 µm; virtual slices from electron tomogram, 200 nm.

### Cmp7 cooperates with SPB proteins to maintain a NE diffusion barrier during mitosis

We next investigated if the loss of fitness of cells lacking Cmp7 and fully functional SPB proteins could be attributed to a specific disruption of a mitotic NE diffusion barrier. To address this, we analyzed the distribution of MGM4 in several genetic backgrounds in which SPB components were perturbed using both temperature sensitive alleles such as *pcp1-15* (Figure 6C) or rapid depletion (1 h after 5IAA addition) in strains expressing AID-fusions of Pcp1 or Ppc89, another core SPB protein (Supplementary Figure S5). We first tested MGM4 localization in *pcp1-15* and *cmp7Δpcp1-15* cells grown at 30°C and after incubation for 3 h at 36°C, the non-permissive temperature for the *pcp1-15* allele (Figure 6C). At 30°C MGM4 was excluded from the nucleus with N/C fluorescence ratios <0.75 in most cells (Figures 6C and 6D). While cells harboring the *pcp1-15* allele that likely transited mitosis at 36°C were slightly more likely than WT cells to display aberrant nuclear MGM4 fluorescence in the subsequent G_1_/S (∼20% of cells had N/C ratios >0.75), the concomitant absence of Cmp7 led to a far more penetrant phenotype (65% of *cmp71¢.pcp1-15* G_1_/S cells had N/C ratios >0.75) (Figure 6C and 6D). We confirmed that the MGM4 entered the nucleus specifically during the preceding mitosis by timelapse imaging of *cmp7Δpcp1-15* cells during a temperature shift from 30°C to 36°C. Strikingly, 91% (N=75) of cells with nuclei that became permeable to the MGM4 reporter during this time course were undergoing mitosis (Figure 6E).

The Cmp7-dependent loss of nucleocytoplasmic compartmentalization in *pcp1-15* cells was largely mirrored by the conditional depletion of either Pcp1-AID or Ppc89-AID. In these cases, however, simply appending the AID protein domain to Pcp1 and Ppc89 sensitized these strains to loss of Cmp7 and resulted in the nuclear entry of MGM4 specifically in *cmp71¢.pcp1-AID* and *cmp71¢.ppc89-AID* G_1_/S cells. Critically, we find that depletion of Pcp1-AID and Ppc89-AID manifests in loss of the NE diffusion barrier (Supplementary Figure S5B, S5C, S5E, and S5F) prior to leading to mitotic SPB and/or spindle defects (Supplementary Figure 5G–5H), although phenotypes such as monopolar spindles do eventually arise in the population (Supplementary Figure 5I). This surprising result suggests that the canonical role of these SPB proteins in organizing the mitotic spindle (Flory et al., 2002; Fong et al., 2010; Rosenberg et al., 2006) may be secondary to their role in maintaining a mitotic NE diffusion barrier.

Lastly, to provide additional insight into the loss of compartmentalization when we combine perturbations of Cmp7 and Pcp1 we performed CLEM of *cmp7Δpcp1-15* cells following 3 h at the restrictive temperature (36°C). We observed that in half (N=6) of the *cmp7Δpcp1-15* cells examined, and none of the *pcp1-15* cells (N=6), there was a failure to exclude ribosomes from the nucleoplasm (Figure 6F and 6G, magnified boxes) in the presence of an otherwise intact NE. Thus, in aggregate our data strongly suggest that maintaining robust nucleocytoplasmic compartmentalization during mitosis requires cooperation between Cmp7 and functional SPB proteins. Moreover, these data further demonstrate that NPCs are not the sole proteinaceous diffusion barriers that function at the *S. pombe* NE.

## DISCUSSION

Here we develop and exploit the fission yeast *S. pombe* as a model of ESCRT function at the NE (Supplementary Figure S6). The data firmly establish that ESCRTs are required to seal the NE hole generated by the extrusion of the SPB. Critically, however, we also propose a new function for ESCRTs prior to sealing in which they act as a molecular grommet that restricts the size of the persistent NE fenestration during anaphase B. This grommet function is needed to support a proteinaceous diffusion barrier that is essential to maintain nucleocytoplasmic compartmentalization. Thus, in addition to extending the functional repertoire of ESCRTs, we also introduce the exciting concept that, beyond the NPC, there are other protein diffusion barriers that restrict the passage of macromolecules across the NE.

Using genetic and functional assays we identify core components of the SPB as essential to a diffusion barrier that acts in concert with the ESCRT grommet at the NE in fission yeast mitosis (Figure 6). While such a function has not been described previously, the well-established concept that the nuclear compartment is defined by NPCs requires that the constitutively inserted SPB in budding yeast (Byers and Goetsch, 1975; O’Toole et al., 1999) is also capable of restricting the diffusion of macromolecules. Taken together, we think it likely that core SPB components across divergent yeasts possess a fundamental physiochemical property that allows them to constitute a diffusion barrier, which intuitively had to evolve coincident with the adoption of closed mitosis.

Our data thus highlight a broader theme, together with published work, in which ESCRTs and proteinaceous diffusion barriers act in concert to protect the integrity of the nuclear compartment. A clear corollary, for example, is how the NPC diffusion barrier installed during NPC biogenesis is supported by the ESCRT pathway in the event that NPC biogenesis fails (Thaller *et al*., 2019; Thaller *et al*., 2021; Webster *et al*., 2014; Webster *et al*., 2016). Another example might include the rapid recruitment of barrier to autointegration factor (BAF) to exposed chromatin at induced NE ruptures (Halfmann *et al*., 2019). As BAF forms a cross-linked mesh network that can restrict the passage of macromolecules 500 kDa/49 nm in size (Samwer et al., 2017), it is possible BAF together with other factors (including lamin C (Kono et al., 2022)), might plug NE holes quickly before the membranes themselves are later sealed through a mechanism involving LEM protein(s) and ESCRTs. A similar function could be served by galectin, which appears capable of forming biomolecular condensates (Chiu et al., 2020) and recruiting ESCRT-III (Jia et al., 2020) to prevent leakage of lysosomal hydrolases during ESCRT-mediated repair of lysosome membranes (Radulovic et al., 2018; Skowyra et al., 2018).

Exactly what physiochemical mechanism(s) underlie the ability of the SPB to form a diffusion barrier remains unclear. One attractive mechanism to form this barrier would be through a biomolecular condensation similar to the canonical diffusion barriers acting at the NE, i.e. the FG-nucleoporins residing in the central channel of NPCs (Celetti et al., 2020; Frey et al., 2006; Petri et al., 2012). Indeed, like NPCs (Ribbeck and Görlich, 2002; Riquelme Barrientos *et al*., 2023; Shulga and Goldfarb, 2003) and many other liquid or hydrogel-like biomolecular condensates (Kroschwald et al., 2017), we find that the diffusion barrier that supports the maintenance of nuclear compartmentalization in late mitosis is sensitive to relatively low concentrations of 1,6-HD (Figure 5). Further, as this diffusion barrier must be able to form over NE holes that span from tens to hundreds of nanometers in diameter (Figure 3), it must possess a plasticity that could be consistent with an unstructured biomolecular condensate. A key candidate that might contribute to such a function may be Pcp1, as its human ortholog, pericentrin, can undergo biomolecular condensation as a component of the pericentriolar material surrounding centrosomes (Feng et al., 2017; Jiang et al., 2021; Woodruff et al., 2017; Zwicker et al., 2014). We note, however, that we previously demonstrated that Pcp1 molecules incorporated into the SPB behave essentially as a solid material, undergoing little to no turnover with the soluble Pcp1 pool (Wälde and King, 2014). Thus, we cannot rule out that the diffusion barrier identified here could also be created through a more structured, mesh-like mechanism, similar to what has been proposed for BAF (Samwer *et al*., 2017). Future work will be needed to explore the physiochemical properties of this barrier more fully and exactly how various proteins, such as those identified here, contribute.

That the diffusion barrier we identify is sensitive to ESCRT perturbation is presumably because larger holes put greater reliance on this proteinaceous plug. Building on this, we propose that ESCRTs act as a molecular grommet that restricts the size of the NE hole during anaphase B to a scale that can be effectively supported by the diffusion barrier. Superficially, this grommet model has similarities to the “O-ring seal” model that has been suggested to reside at sites of reforming NE surrounding kinetochore-attached microtubules in mammalian cells (von Appen *et al*., 2020). The concept of an O-ring is primarily based on the formation of a ring-shaped CHMP7-LEM2 co-polymer that is observed surrounding microtubules *in vitro* (von Appen *et al*., 2020)— the structure formed *in vivo* remains unknown. However, our quantitative analysis reveals that only ∼16 molecules of Cmp7 reside at the NE hole in anaphase B (Table 1 and Supplementary Figure S2), far less than would be necessary to line the 50–150 nm NE hole (corresponding to the widths measured when Cmp7 is recruited to the SPB extrusion site; Figure 3) if it forms a co-polymer analogous to that suggested for the mammalian CHMP7-LEM2 (von Appen *et al*., 2020). We suggest that the grommet model outlined here is distinct, favoring instead the possibility that Cmp7 (the sequence of which contains elements of both ESCRT-II and ESCRT-III family members) is a seed filament for the assembly of additional downstream ESCRTs, akin to models that suggest a sequential assembly order of ESCRT-IIIs at endosomes (Babst et al., 2002; Henne et al., 2012; Pfitzner et al., 2020; Saksena et al., 2009; Teis et al., 2008) as well as other contexts that begin with ESCRT-I and ESCRT-II proteins (Vietri *et al*., 2020). Such a model predicts that Cmp7 is required for all ESCRT recruitment to the NE, which is indeed the case (Figure 2). As CHMP7 is also required for ESCRT subunit recruitment to the NE in mammalian cells at mitotic exit (Gu *et al*., 2017; Olmos *et al*., 2016), we suggest that the fundamentals of ESCRT recruitment and function in *S. pombe* and mammalian cells are highly similar. Subtleties between the two models will need to be revealed in future studies that examine the copy number of ESCRTs at well-defined NE holes in mammalian cells in the manner that we have performed here.

Although the observed copy number of Cmp7 molecules indicates that they may not be able to independently line a 50–150 nm NE hole, they may nonetheless be constituents of a more complex polymer that includes the Wave 2 ESCRTs, Vps32 and Ist1, which are found at much higher copy numbers than Cmp7 (Table 1 and Supplementary Figure S2). Indeed, based on the recruitment timing (Figure 1) and its relationship to the normal constriction of the NE hole (Figure 3), we infer that a polymer(s) consisting of Wave 1 and 2 ESCRTs comprises the grommet. This grommet appears to restrict the NE hole diameter to support the SPB protein-dependent diffusion barrier (Figure 6). However, as we observe that the NE hole shrinks from 150 nm to 50 nm during the Wave 1 and 2 recruitment period, it is also possible that the grommet is responsible for physically driving its constriction. We speculate that this function could be attributed to Ist1, which is thought to contribute to ILV neck constriction prior to membrane scission during ILV formation (McCullough et al., 2015; Nguyen et al., 2020; Pfitzner *et al*., 2020), with the caveat that the dimensions of a constricting ILV is an order of magnitude smaller than the NE hole. This comparison further highlights a key difference between the order of assembly/composition of ESCRTs at the NE compared to at endosomes, where Ist1 is instead recruited much later than Vps32 (Pfitzner *et al*., 2020). Ultimately, a model of membrane fission at the NE must incorporate this difference, as well as the apparent absence of the ESCRT-III Did2; we note, however, that the mammalian Did2 ortholog, CHMP2A, was reported to localize to the reforming NE at mitotic exit (Vietri *et al*., 2015), opening up the alternative possibility that the levels of Did2-GFP present are below our threshold of detection (Supplementary Figures S1A and S1B).

Our data clearly underscore that ESCRTs are required for the penultimate step of NE sealing, likely through a membrane fission mechanism (Figure 3). However, our observation of NE holes as small as 20 nm in the absence of Cmp7 suggests ESCRT-independent mechanisms capable of minimizing the persistent NE hole in G_2_, possibly involving local lipid synthesis or flow of membrane from the ER (Kinugasa et al., 2019; Kume et al., 2019; Penfield *et al*., 2020); normal nucleocytoplasmic compartmentalization in this context further suggests the presence of a functional barrier to diffusion residing in the persistent NE hole. Interestingly, our data also support that the NE sealing mechanism requires maturation of the ESCRT filament including a wholesale replacement from a Cmp7-Vps32-Ist1 grommet in late anaphase B to a NE sealing modality consisting of Vps24 and Did4 in the subsequent G_1_/S (Figures 2 and 4). Indeed, temporally, the release of Cmp7 occurs in concert with recruitment of the Wave 3 ESCRTs (Figure 1). We postulate that Cmp7 acts as a seed that initiates a sequence of ESCRT recruitment events, ultimately leading to a mature state of a Wave 3 ESCRT filament that drives NE sealing. Disrupting Wave 3 ESCRTs (Vps24 or Did4) drives the persistence of a Wave 1 state in which Cmp7 is found at an abnormally high copy number, apparently stuck in the NE hole that fails to seal (Figures 2 and 4).

The observations that both disrupting ESCRT recruitment (*cmp7Δ* cells, Figure 3) or accumulating a stalled ESCRT assembly (*did4Δ* cells, Figure 4) compromise NE sealing has several implications for interpreting how loss of function of ESCRTs manifest more generally. As nucleocytoplasmic compartmentalization can appear grossly normal despite the persistence of these small NE holes, their presence may be undetected by monitoring reporter proteins (Figure 5A). Thus, these observations highlight that it will be important to revisit data suggesting ESCRT-independent NE sealing mechanisms that did not include ultrastructural analysis. Perhaps most importantly, we think it likely that persistent NE holes could lead to an unstable NE barrier that is more susceptible to further perturbation. As loss of Cmp7 and persistent Cmp7 converge in precipitating a failure of NE sealing, this work provides a useful framework to interpret the consequences of ESCRT loss or gain of function in the context of disease. For example, variants in the CHMP2B gene (ortholog of Did4) cause frontotemporal dementia (Skibinski et al., 2005) and the aberrant nuclear entry of CHMP7 is upstream of a pathological cascade that could contribute to both sporadic and familial forms of amyotrophic lateral sclerosis (Coyne et al., 2021). In both scenarios, it is possible that improper clearance of CHMP7 could compromise the integrity of the NE over timescales beyond that easily monitored in cell culture. Further, it will be interesting to examine if some cancer cells are prone to repeated nuclear ruptures due to NE nanoholes arising from inefficient ESCRT-mediated repair, rendering them more fragile.

## ACKNOWLEDGEMENTS

We thank the Center for Cellular and Molecular Imaging, Electron Microscopy Facility at Yale School of Medicine for assistance with EM, particularly Morven Graham and Xinran Liu. Additionally, we thank Li-Lin Du, Adam Frost, Iain Hagan, Tom Pollard, and NBRP (YGRC), Japan for yeast strains. This work was supported by the National Institutes of Health (R01 GM105672 to C.P. Lusk; F32 GM139285 to N.R. Ader) and an Allen Distinguished Investigator Award, a Paul G. Allen Frontiers Group advised grant of the Paul G. Allen Family Foundation (M.C. King).

## METHODS

### *S. pombe* culture and strain generation

Strains used in this study are listed in Supplemental Table S2. Strains were cultured and mated using standard procedures and media (Moreno et al., 1991). In brief, *S. pombe* were cultured on yeast extract supplemented with 250 mg/liter adenine, histidine, leucine, uracil, and lysine hydrochloride (YE5S) agar plates and liquid cultures. For experiments with MGM4, cells were grown in Edinburgh minimal medium supplemented with 250 mg/liter adenine, histidine, leucine, uracil, and lysine hydrochloride (EMM5S) overnight prior to imaging. C-terminal tagging and gene replacement (knockout) were performed using the pFA6a, MX6-based drug resistance cassettes (Bahler et al., 1998; Hentges et al., 2005) using *S. pombe* genome information available through PomBase (Harris et al., 2022; Wood et al., 2002). Visualization and virtual manipulation of DNA sequences was facilitated by using ApE 2.0.61 (Davis and Jorgensen, 2022). For all GFP fusions, mEGFP was used. For C-terminal tagging of Cmp7 for AID depletion we used pNA55 (pDB4581), a gift from Li-Lin Du (Addgene plasmid # 171124) (Zhang *et al*., 2022). All genomic integrations were confirmed by screening of individual clones by PCR of whole yeast (colony PCR) (Huxley et al., 1990) with primers designed to amplify only a genomic insertion at the targeted locus following a standard lithium acetate transformation procedure (Murray et al., 2016).

As has been previously shown, dysfunctional tagged ESCRT proteins display a loss of dynamic remodeling and aberrantly accumulate in class E endosomes (Teis *et al*., 2008). Thus, to generate functional GFP-ESCRTs, we first attempted C-terminal tagging using a 3×HA linker. For Cmp7-3×HA-GFP, Ist1-3×HA-GFP, and Did2-3×HA-GFP this strategy yielded GFP-ESCRTs which were dynamic by live-cell imaging (Figure 1) and capable of generating progeny when mated. As we noted that expression of Vps32-3×HA-GFP and Did4-3×HA-GFP appeared to drive a dominant negative phenotype that inhibited tetrad formation, we used an N-terminal tagging strategy for tagging of Vps32, Did4, and Vps24. N-terminal GFP-tagging of Vps32, Vps24, and Did4 was performed by inserting a pJK148 plasmid cut with NruI at the *leu1-32* locus in a strain with the endogenous gene knocked out (pNA32 for GFP-3×HA-Vsp32, pNA23 for GFP-3×HA-Vps32, and pNA45 for GFP-3×HA-Did4; see Supplementary Table S2 for full plasmid list) (Keeney and Boeke, 1994). To generate pNA22, pNA23, and pNA45 plasmids, we performed Gibson Assembly of the following sequences: (1) promoter (500 bp upstream of start 5’ UTR) + 5’UTR, (2) GFP-3×HA, (3) open reading frame + 3’UTR + terminator (500 bp downstream of stop codon) (Gibson et al., 2009). Strains with GFP-3×HA -Vps32, GFP-3×HA -Vps24, and GFP-3×HA -Did4 had dynamic GFP-ESCRT foci (Figure 1) and had no apparent defect in mating, producing viable tetrads.

### Live-cell fluorescence microscopy

Cells expressing fluorescent protein fusions were grown to log phase (OD_600_ 0.4–0.8) in YE5S supplemented with an additional 250 mg/L adenine (to ensure low autofluorescence) and 0.1 mM *n*-propyl gallate (NPG) (an oxygen scavenger to reduce photobleaching and mitochondrial autofluorescence), concentrated by 20 s centrifugation at 2000× *g*, sandwiched between a #1.5 glass coverslip (VWR 48393) and a glass slide (Thermo Fischer Scientific 420), and sealed with VALAP (1:1:1, vaseline:lanonin:paraffin). Image acquisition was performed on a DeltaVision microscope (Applied Precision; GE Healthcare) using Resolve3D in SoftWoRx using the Insight SSI 4-color Live Cell filter set (ex425–495/em500–550 nm for GFP, ex555–590/em600–675 nm for mCherry) on a CoolSNAP HQ^2^ CCD (Photometrics) or Evolve-512 EMMCD (Photometrics) with either an UPlanSapo 100× 1.4 NA oil immersion objective (Olympus) or a PlanApo 63× 1.42 NA oil immersion objective (Olympus) and an AURA light engine (Lumencor) at 32% transmission or 100% transmission for up to 1 s. Cells were imaged over 4.8–5 µm in the z-plane at 0.2 intervals for counting, 0.5 µm for compartmentalization (MGM4 and MGM2), and 0.2–0.4 µm for all other experiments. Deconvolution was performed using SoftWoRx; deconvolved images were used only for display, not intensity based analysis. For timelapse experiments with drug delivery, cells were first loaded into a CellASIC ONIX microfluidic plate (Millipore Y04C) before imaging. Microfluidics were controlled by the CellASIC ONIX microfluidic platform (Millipore EV262, MIC230, GM230). For Cmp7-AID depletion, 5IAA (TCI Chemicals, A3390) was dissolved in DMSO to make a 1 mM stock solution and used at a final concentration of 100 nM in culture. For treatment with 1,6-HD, 1,6-hexanediol (Alfa Aesar, A12439) was dissolved in YE5S plus 250 mg/L adenine to make a 25% stock solution, which was diluted to either 2.5% or 5% as needed. Fluorescence micrographs were viewed and analyzed using Fiji 2.3.0/1.53q unless otherwise noted (Schindelin et al., 2012).

### Image Display

All representative micrographs have been deconvolved as described above and are displayed to maximize the displayed dynamic range without pixel saturation in the linear range. When mCherry-Atb2 and Pcp1-mCherry or mCherry-Atb2 and Sad1-mCherry are displayed together, the gamma has been adjusted to 0.05 to allow both fluorophores to be displayed without pixel saturation. All magnified ROIs of fluorescence micrographs were independently adjusted.

### Cell cycle classification for GFP-ESCRT localization to SPB extrusion site

To classify cells into phases of the cell cycle from still images, we used the following criteria: Cells with mitotic spindles longer than 3 µm were considered to be in anaphase B and were converted to time in anaphase B using the following formula:

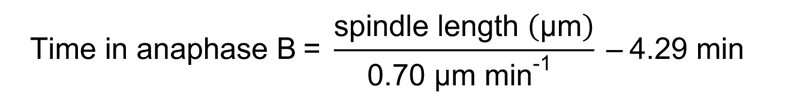

We used the rate of spindle elongation of 0.7 µm/min in anaphase B as determined in (Krüger *et al*., 2019) and the correction factor of 4.29 min represents a spindle length of 3 µm at the anaphase A/B transition. Cells with mitotic spindles longer than 3 µm, but smaller than 6.35 µm were classified as early anaphase B (approximately 0–5 min into anaphase B). Cells with spindles between 6.35 µm and 8.39 µm were classified as mid anaphase B (approximately 5–7.5 min into anaphase B) and any cell with a spindle longer than 8.39 µm was classified as late anaphase B (>7.5 min into anaphase B). Late G_1_/S cells were distinguished from early G_1_/S cells by the lack of microtubules spanning the septum region. For each phase of the cell cycle the percentage of SPB foci (Sad1-mCherry or Pcp1-mCherry) that colocalized with the GFP-ESCRT was quantified for still images as shown in Figures 1C, 2C–E, 2G, and 2I as well as Supplementary Figures S1B and S3C. Quantification was performed across three biological replicates with at least 50 nuclei per replicate.

### Assessing the timing of GFP-ESCRT at SPB in anaphase B

Strains expressing a GFP-ESCRT protein, a SPB marker (Sad1-mCherry or Pcp1-mCherry), and mCherry-Atb2 were imaged every 3 min for approximately 20 min. For cells entering anaphase B, we recorded the length of the mitotic spindle in the first frame of the time-series where a focus of the GFP-ESCRT could be discerned. These spindle lengths were converted to time in anaphase B using the above formula (Figure 1D). We also used this method to quantify the timing of SPB extrusion and NE constriction (Figure 1D, vertical gray bars). Quantification was performed across three biological replicates with at least 50 nuclei per replicate.

### Quantification of average copy number of ESCRT-III subunits

Cells were prepared for imaging by overnight culture in YE5S plus 250 mg/L adenine + 0.1 mM NPG to log phase. Cells (MKSP3186) expressing Fta3-GFP were mounted on the same pad as a GFP-ESCRT-expressing strain of interest. Cells were imaged as above. Raw images were then analyzed using a custom MATLAB script, ‘Find_n_Fit_Spots’ (https://github.com/LusKingLab/Find_n_Fit_Spots_3D). First, spots were detected using ‘findIrregularSpots3’, a custom-written MATLAB code that finds fluorescent objects regardless of their shape. Briefly, the code identifies all the brightest pixels within sub-volumes of a given size. Then, it calculates the ratio of the mean intensities of “central” pixels over “edge” pixels. If the ratio is above a given threshold, the code keeps the identified spot and calculates its 3D coordinates as an intensity-weighted centroid of pixels. Second, for each “good” spot, x-y-coordinates were used to crop a circular image with R = 7 pixels around the spot from the best 2D image (determined by z-coordinate) which was fitted by the linear combination of a 2D Gaussian and error function, *erf*, to account for nuclear edge background signal. The volume of the best Gaussian fit was then used as a spot intensity. Identified spots were then manually curated to classify the spot (Fta3-GFP or GFP-ESCRT) based on secondary fluorescent markers. Position in the cell cycle was determined using brightfield images and, for GFP-ESCRT strains, mCherry-Atb2. For each GFP-ESCRT focus, the copy number was determined by dividing the intensity of the focus by the average intensity of mitotic Fta3-GFP, previously calculated to be approximately 111 molecules (Lawrimore *et al*., 2011). For the comparison of Fta3-GFP intensities across cell cycles (Supplemental Figures S2E–S2G), the measured intensity is plotted by position in the cell cycle. To allow comparisons across three biological replicates, individual datapoints were normalized to the average value for each cell cycle position within the replicate. Fta3-GFP or GFP-ESCRT foci identified as outliers using ROUT test Q = 1%, 1 iteration were excluded.

### Quantification of N/C ratio of fluorescent reporters

To calculate the ratio of MGM4 and MGM2 fluorescent reporters in the in nucleoplasm vs. cytoplasm, we measured the mean intensity of regions within the nucleoplasm, cytoplasm (avoiding vacuoles), and outside of the cells (background). The background mean intensity was subtracted from both the nucleoplasm and cytoplasmic mean intensities before calculating the N/C ratio. The individual N/C ratios were then binned with 0.05 bin size using Microsoft Excel. When these data are displayed as a histogram (Figure 5D–G), where the percentage of nuclei in each bin is shown. For visual simplicity, these data are simplified and displayed as only the percent of cells with nuclear MGM4 (nuclei in bins 0.75 and above; Figure 6D and Supplementary Figures S5C and S5F). For percentage change of N/C ratio in time-series imaging (Figure 5I), the N/C ratio of the first frame of the movie was subtracted from the N/C ratio of the last frame and then divided by the N/C ratio of the first frame. Quantification was performed across three biological replicates with at least 50 cells per replicate.

### Quantification of MGM4 nuclear fluorescence over time

To analyze the fluorescence intensity of nuclear MGM4 reporter, we first corrected for photobleaching using the “Bleach Correction” function in Fiji with an exponential fit. For each cell, we measured the mean fluorescence intensity of a region within the nucleoplasm for each frame of a time series. To compare cells within and across populations, background subtracted intensity measurements were normalized to the intensity of the nuclear region in the first frame. Quantification was performed across three biological replicates with at least 5 nuclei per replicate.

### Quantification of Cmp7-GFP fluorescence intensity in *did4Δ* cells

To compare the intensity of Cmp7-GFP at the SPB site across the cell cycle, we measured the mean intensity the GFP channel in a background-subtracted circular area. Position in the cell cycle was determined by mCherry-Atb2 morphology. To allow comparisons across three biological replicates, individual datapoints were normalized to the average value for mitotic cells with visible Cmp7-GFP foci. At least 30 cells were analyzed in each of three biological replicates.

### Correlative light and electron microscopy

CLEM of resin-embedded cells was performed as described by Kukulski et al. (2012). In brief, cells were grown overnight in YE5S plus 250 mg/L adenine to log phase before being concentrated using either filtration, with a 0.45 µm nitrocellulose membrane (Millipore HAWP02500) and a microanalysis filter setup (Millipore XX1012500) (Figure 3), or centrifugation at 2000× *g* for 20 s (Figure 6F, G). Concentrated cells were transferred using either a blunted toothpick or micropipette to the 200-µm recess of an aluminum platelet (Engineering Office M. Wohlwend 241) (McDonald, 1999). Platelets were then frozen using a high-pressure freezer (Leica Microsystems HPM100). Samples were freeze substituted in 0.1– 5% uranyl acetate in acetone and embedded in Lowicryl HM20 (Polysciences) with automated temperature control (Leica Microsystems EM-AFS2), manual agitation and solution exchange. Resin-embedded cells were sectioned to a 250 nm nominal thickness using an ultramicrotome (Leica Artos 3D) equipped with a diamond knife (Diatome) before being collected on 200 mesh copper grids with carbon support (Ted Pella 01840).

Fluorescence micrographs of grids were acquired using a DeltaVision microscope with the optical configuration as described above for live-cell imaging. For each grid square of interest, 10 z-sections were acquired every 250 nm with 3 s exposure with GFP and mCherry imaging setup at 32% transmission. 15 nm protein A-coated gold beads (Cell Microscopy Core, University Medical Center Utrecht) were adhered to grids prior to EM to aid in tilt-series alignment prior to tomogram reconstruction.

Single or dual-axis tilt series were collected on an electron microscope (FEI TF20) operated at 200 kV using a high-tilt tomography holder (Fischione Instruments 2020) from approximately -60 to +60 degrees (one-degree increments) at a binned pixel size of 1.242 nm on a 4k × 4k Eagle CCD (FEI) using a 150 µm C2 aperture and a 100 µm objective aperture in an automated fashion using SerialEM (Mastronarde, 2005). Reconstruction and segmentation were performed using IMOD (Kremer et al., 1996), with the former in an automated fashion (Mastronarde and Held, 2017). For better visibility in shown virtual slices, a nonlinear anisotropic diffusion and Gaussian filter were applied in IMOD. Fluorescence micrographs have been rotated to match the orientation of virtual slices and adjusted for contrast individually.

### Measurement of NE hole width

To measure the size of the NE hole at the SPB extrusion site, we recorded the maximum 2D distance between the two pore membrane regions of the NE abutting the SPB through the tomogram in IMOD. As some tomograms captured only part of the NE in the visualized slice, some of these values may be underestimates of the actual hole size (Figure 3I, open circles).

### Preparation of whole cell protein extracts and immunoblotting

Overnight cultures of yeast were grown in EMM5S to log phase. Cultures were centrifuged at 2000× *g* for 5 min and cell pellets were washed with 1 mM EDTA. Cells were then pelleted and lysed in 2M NaOH incubated for 10 min on ice. An equal volume of 50% trichloroacetic acid (TCA) was mixed in before incubating for 10 min on ice and collecting the protein precipitate by centrifugation. The pellet was then washed with –20°C acetone and air-dried for 15 min. The pellet was then dissolved in 5% SDS followed by an equal volume of SDS-PAGE sample buffer containing urea (24 mM Tris-Cl pH 6.8, 9 M urea, 1 mM EDTA, 1% SDS, 10% glycerol). Samples were either shaken at 37°C for 15 min (Supplementary Figure S3B) or incubated at 90°C for 5 min (Supplementary Figures S5A and S5D) before being centrifuged at 14,000× *g* for 15 min. Approximately equal loads of extracted protein was then resolved on tris-glycine 10% SDS-PAGE gels and transferred using wet transfer to 0.2 µm nitrocellulose membrane (Bio-Rad 1620112). Transferred proteins were then stained with Ponceau S Solution (Sigma-Aldrich P3504) before being scanned on a flatbed scanner (Epson V800 Photo). Blots were blocked overnight in blocking buffer (5% (w/v) dry milk/TBST) and incubated with primary mouse α-mAID antibody (MBL M214-3) (diluted 1:1000 in blocking buffer). The blots were then incubated for 1 h at room temperature with secondary HRP-conjugated goat anti-mouse antibodies (Invitrogen 3430). Antibody-labeled proteins were visualized using SuperSignal West Femto Maximum Sensitivity ECL Substrate (Thermo Fischer Scientific 34094) on a VersaDoc Imaging System (Bio-Rad 4000 MP) with a 1.4 mm aperture and a three-minute exposure time. Digital images of blots were processed using the “Despeckle” function in Fiji and then adjusted to maximize intensity range without over-or under-saturation.

### Plate-based growth assays

To evaluate relative colony growth rates, cells were grown overnight in YE5S media. Cells were then diluted to 1 OD_600_ equivalent in YE5S before sequentially plating 5 serial 1:10 dilutions on YE5S plates. The plates were incubated at the indicated temperatures in figures/figure legends and scanned daily on a flatbed scanner (Epson V800 Photo).

### Statistical methods

All graphs were generated and analyzed for statistical significance using Prism 9 (GraphPad). Tests used for each figure panel are denoted in figure legends.

### Data Availability

The datasets generated during and/or analyzed during the current study are available from the corresponding authors on reasonable request.

### Code Availability

MATLAB script for quantification of ESCRT subunits is available here with no access restrictions: https://github.com/LusKingLab/Find_n_Fit_Spots_3D

## SUPPLEMENTARY FIGURES

**Supplementary Figure S1.**
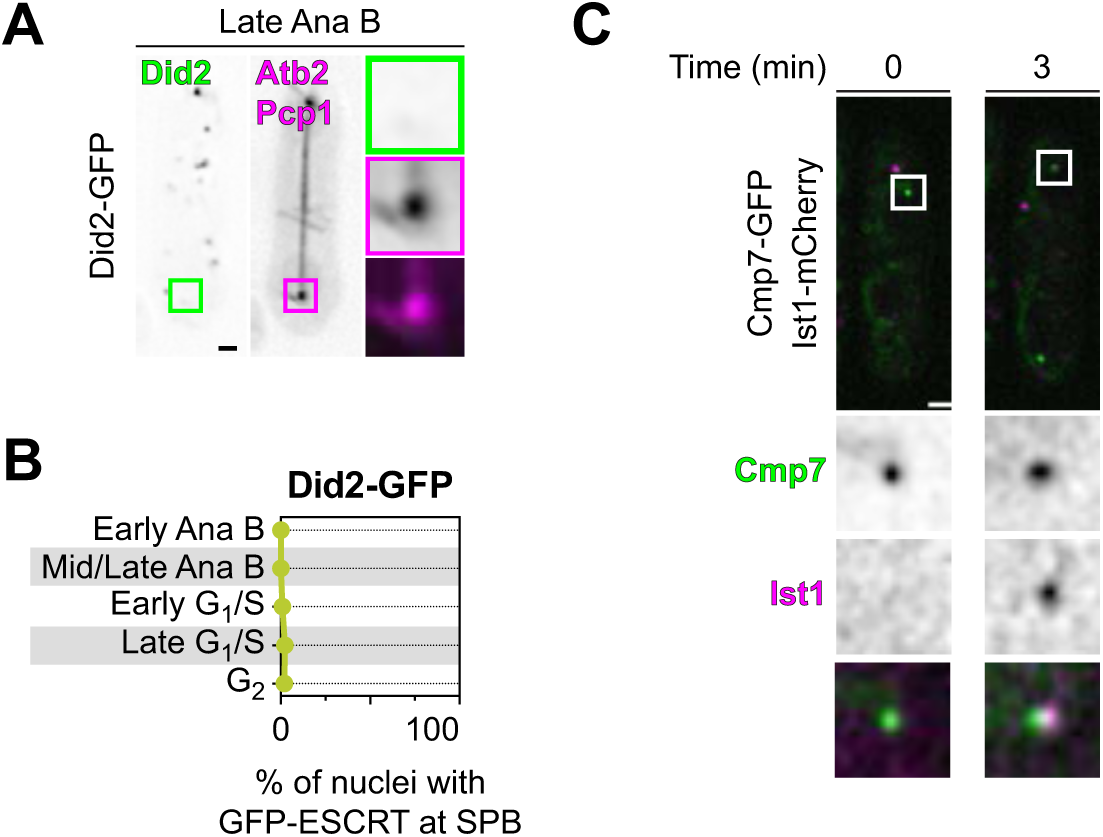
Did2 is not detected at the SPB extrusion site, while Ist1 is recruited after Cmp7. **A:** Fluorescence micrographs of Did2-GFP, mCherry-Atb2, and Pcp1-mCherry in a representative cell in anaphase B. Image of the Cmp7-GFP is on left, sum projection (over 4.8 µm) of mCherry-Atb2 in middle and magnifications of square ROI shown vertically arranged at right with (top to bottom) green, magenta and merged channels. Fluorescence intensity of magnified ROIs were independently adjusted in linear range. Scale bar, 1 µm. Magnified ROIs, 1.6 µm wide. To maximize image clarity, fluorescence intensity of magnified ROIs were independently adjusted in the linear range with the exception of the magenta channel where the gamma was adjusted to 0.05 to allow visualization of mCherry-Atb2 and Pcp1-mCherry without saturation. **B:** Plot of the percentage of nuclei with Did2-GFP (MKSP3342) colocalized with SPB marker at indicated phases of the cell cycle. Points and error bars represent mean and range, respectively, across at least three biological replicates with at least 50 nuclei/GFP-ESCRT/replicate, N = 372 nuclei. **C:** Time-lapse fluorescence microscopy of Cmp7-GFP (green) and Ist1-mCherry (magenta) in a late anaphase B cell (MKSP3521). Cell representative of population from two biological replicates. White boxes in overview micrographs indicate ROI magnified below images as single channel and composite images.

**Supplementary Figure S2.**
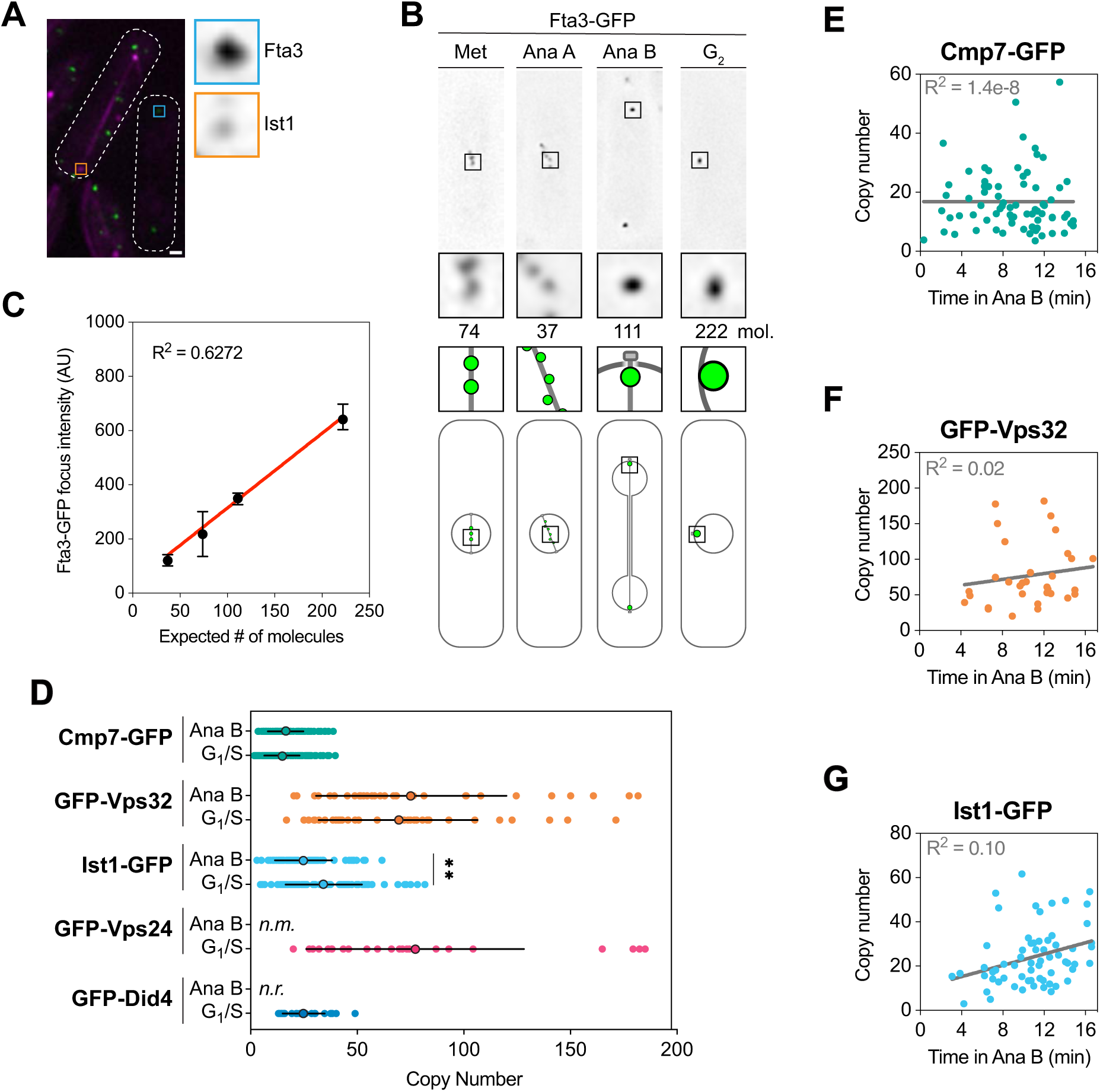
Average copy number of ESCRTs at the SPB extrusion site determined by ratiometric quantification. **A:** Fluorescence micrographs of representative mixture of a GFP-ESCRT strain (Ist1-GFP, MKSP3365) and one expressing Fta3-GFP (MKSP3186). In top left, strain expressing Ist1-GFP (green), mCherry-Atb2 (magenta), and Sad1-mCherry (magenta). In bottom right, Fta3-GFP strain (green). Scale bar, 1 µm. Blue and orange boxes in overview micrographs indicate ROI magnified to the right of image as single channel images for Fta3-GFP and Ist1-GFP, respectively. **B:** Fluorescence micrographs of representative Fta3-GFP expressing cells (MKSP3186) at indicated cell cycle stage (top). Black boxes in overview micrographs indicate ROI magnified below images (middle). Schematic representation of Fta3-GFP appearance by cell cycle show (bottom). Number of molecules based on Lawrimore et al. (2011). **C:** XY plot of Fta3-GFP focus fluorescence intensity against expected number of molecules per focus in MKSP3186 based on Lawrimore et al. (2011). Points represent median of all values collected for that stage of the cell cycle (B); error bars represent 95% confidence interval. Red line represents a simple linear regression with the R^2^ shown. N = 366 foci across three biological replicates with at least 50 foci/replicate. **D:** Number of molecules of indicated GFP-ESCRT at the SPB extrusion site during all phases of the cell cycle. Outlined point and black lines represent mean and standard deviation, respectively. *n.r.*, not recruited. *n.m.*, not measured. Data was collected over at least three biological replicates. Kruskal-Wallis test performed across all cell cycle phases for each ESCRT individually with Dunn’s multiple comparisons across all other cell cycle phases (*, p < 0.05; **, p<0.01; all others, p ≥ 0.05). Fta3-GFP (MKSP3186); Cmp7-GFP (MKSP3363), N = 196 foci; GFP-Vps32 (MKSP3353), N = 68 foci; Ist1-GFP (MKSP3365), N = 147 foci; GFP-Vps24 (MKSP3352), N = 24 foci; GFP-Did4 (MKSP3334), N = 26 foci. **E–G:** Copy number of indicated GFP-ESCRT shown as a function of time in anaphase B. Gray lines represent a simple linear regression with the R^2^ shown in gray. Fta3-GFP (MKSP3186); Cmp7-GFP (MKSP3363), N = 73 foci; GFP-Vps32 (MKSP3353), N = 32 foci; Ist1-GFP (MKSP3365), N = 64 foci.

**Supplementary Figure S3.**
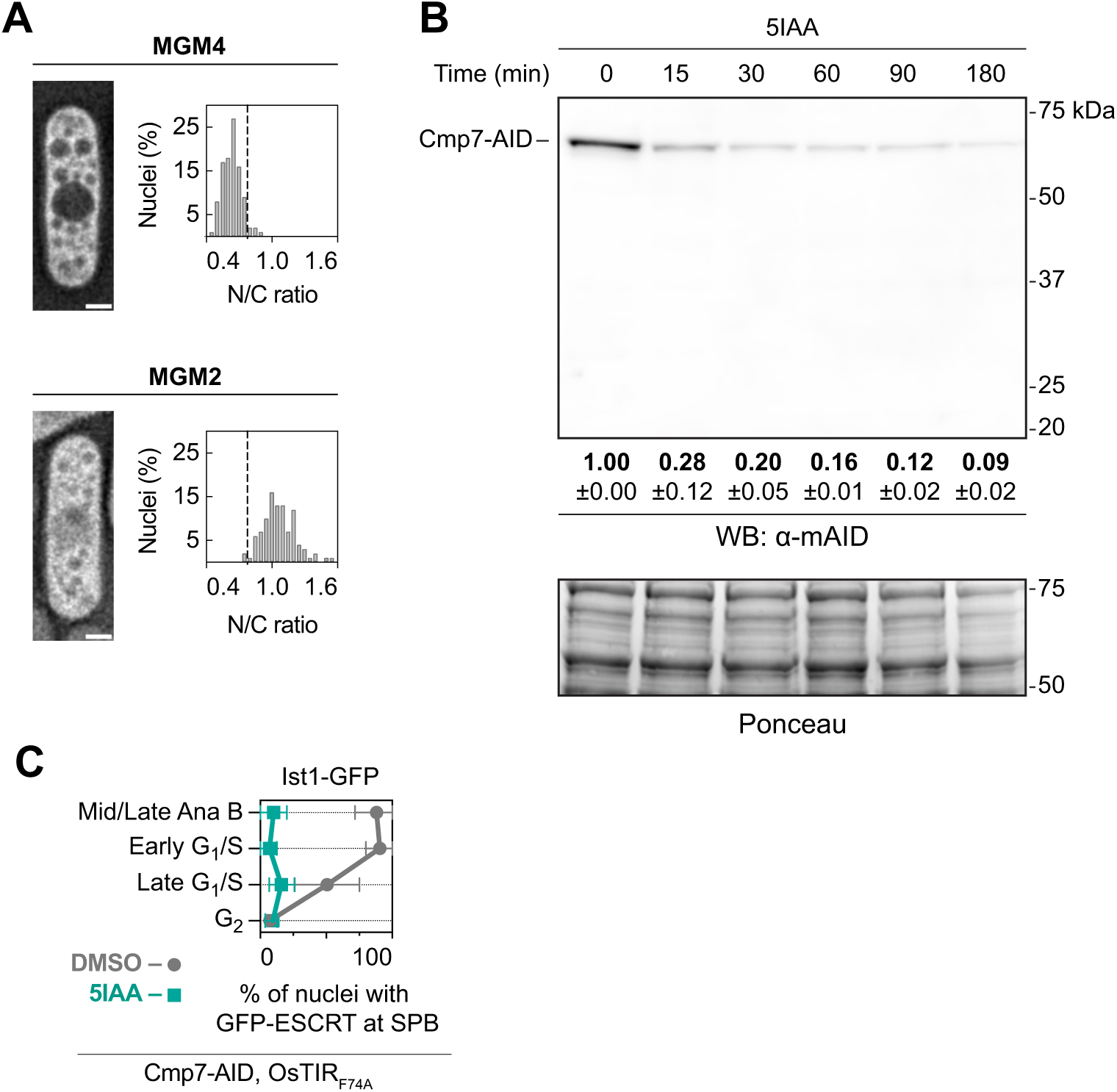
MGM4 and MGM2 reporters; auxin induced degron (AID) system allows for fast degradation of Cmp7. **A:** Deconvolved live-cell fluorescence microscopy of MGM4 (top left, MKSP3345) and MGM2 (bottom left, MKSP3960). Histogram of nucleoplasmic to cytoplasmic GFP fluorescence (N/C ratio) for individual cells shown to the right of each reporter. MGM4 data replotted from Figure 5A. Size between bins, 0.05. Dotted line at 0.75 bin. Quantification was performed for three biological replicates. MGM2, N = 178 nuclei, with at least 58 nuclei/biological replicate. **B:** Representative western blot (WB) of Cmp7-AID levels (detected with an α-AID antibody) in total protein extracts from cells (MKSP3517) grown in the presence of 5IAA for the indicated times. Ponceau stain is shown at bottom to assess relative total protein loads. Relative amount of Cmp7-AID protein, normalized to T0, is shown below WB as mean and standard deviation for three biological replicates. **C:** Plots of the percentage of nuclei with Ist1-GFP colocalized with SPB marker at indicated phases of the cell cycle in cells (MKSP3625) expressing Cmp7-AID and OsTIR1F74A. Cells were treated with DMSO (gray) or 5IAA (green) for 1 h. Points and error bars represent mean and range, respectively, across at least three biological replicates with at least 50 nuclei/GFP-ESCRT/replicate. DMSO, N = 273 nuclei; 5IAA, N = 296 nuclei.

**Supplementary Figure S4.**
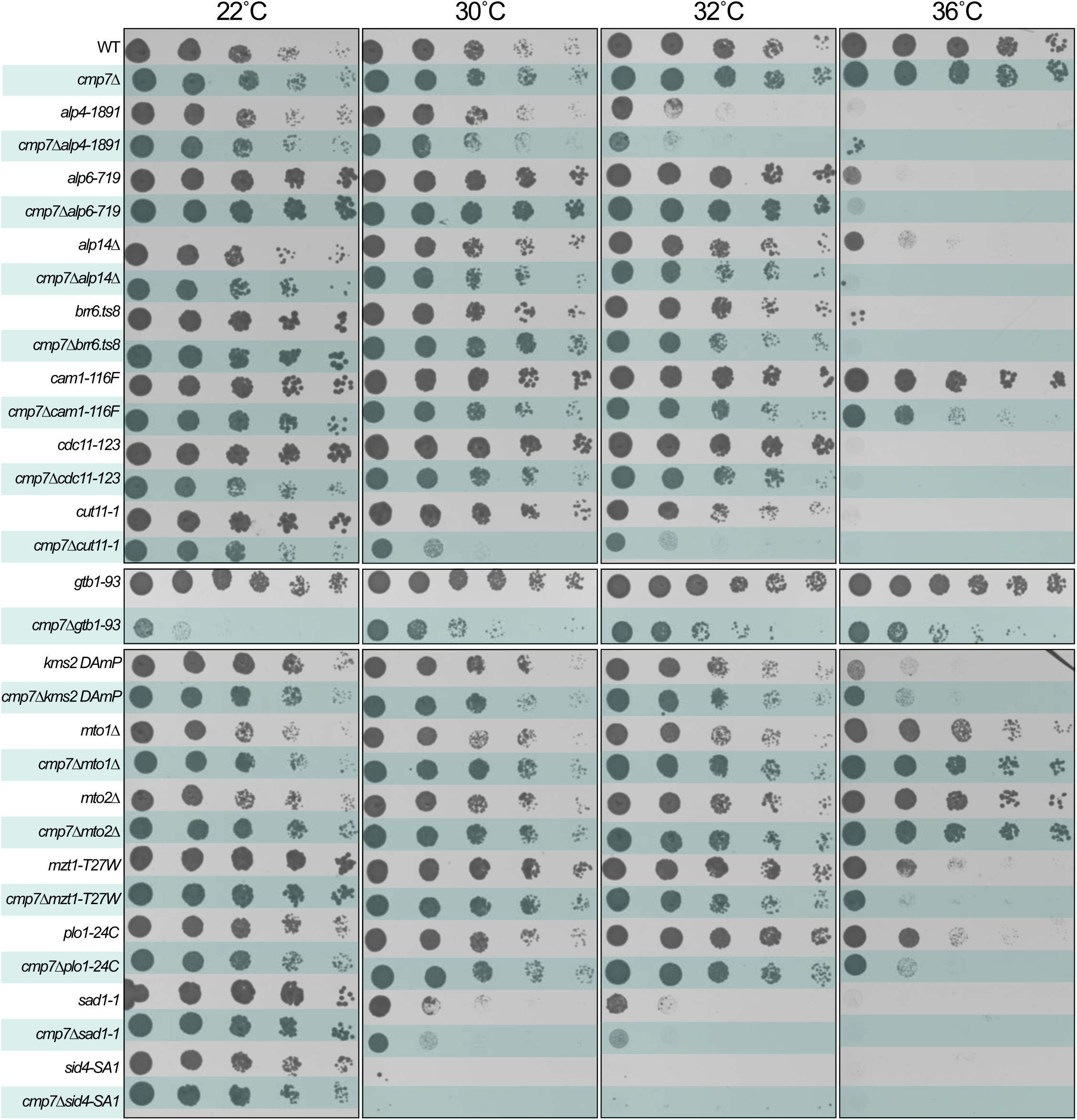
Genes encoding constitutive SPB proteins and mitotically associated SPB proteins genetically interact with *CMP7*. The indicated strains were serially diluted 10-fold and plated on YE5S plates incubated at the indicated temperatures. Plates were imaged after 2 days (30, 32, and 36°C) or 4 days (22°C). WT, MKSP399; *cmp7Δ*, MKSP3107; *cmp7alp4-1891*, MKSP4067; *7Δalp4-1891*, MKSP4163; *alp6-719*, MKSP3475; *cmp7Δalp6-719*, MKSP4031; *alp14Δ*, MKSP4063; *cmp7Δalp14Δ*, MKSP4165; *brr6.ts8*, MKSP3461; *cmp7Δbrr6.ts8*, MKSP3528; *cam1-116F*, MKSP4061; *cmp7*Δ*cam1-116F*, MKSP4167; *cdc11-123*, MKSP4060; *cmp7Δcdc11-123*, MKSP4169; *cut11-1*, MKSP3469; *cmp7Δcut11-1*, MKSP4040; *gtb1-93*, MKSP4062; *cmp7Δgtb1-93*, MKSP4174; *kms2 DAmP*, MKSP1474; *cmp7Δkms2 DAmP*, MKSP3683; *mto1Δ*, MKSP1303; *7Δmto1Δ*, MKSP4027; *mto2Δ*, MKSP1305; *cmp7Δmto2Δ*, MKSP4151; *mzt1-T27W*, MKSP3476; *cmp7Δmzt1-T27W*, MKSP4045; *plo1-24C*, MKSP1461; *cmp7Δplo1-24C*, MKSP4048; *sad1-1*, MKSP3468; *cmp7Δsad1-1*, MKSP4033; *sid4-SA1* MKSP4065; *cmp7Δsid4-SA1*, MKSP4171.

**Supplementary Figure S5.**
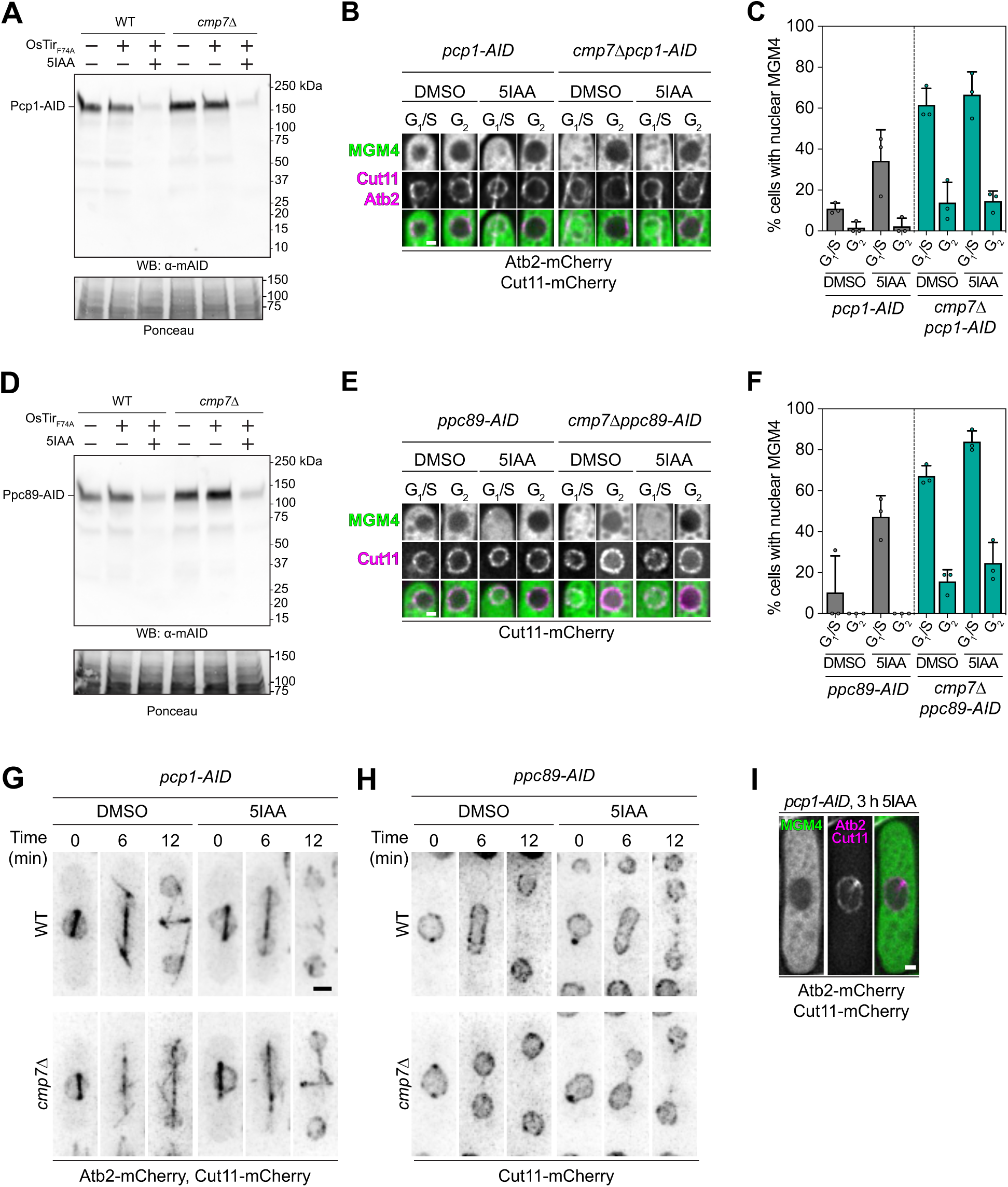
Conditional degradation of Pcp1 and Ppc89 impairs nucleocytoplasmic compartmentalization without impacting mitosis progression. **A:** Western blot (WB) of Pcp1-AID levels (detected with an α-AID antibody) in total protein extracts from cells expressing OsTir1F74A and treated with 5IAA as indicated. Ponceau stain is shown at bottom to assess relative total protein loads. (-)OsTir1F74A (MKSP4112); (+)OsTir1F74A (MKSP4113); (-)OsTir1F74A, *cmp7Δ* (MKSP4114); (+)OsTir1F74A, *cmp7Δ* (MKSP4115). **B:** Fluorescence micrographs of representative nuclei are shown arranged vertically with MGM4 (green) at top, Cut11-mCherry (magenta) and mCherry-Atb2 (magenta) in middle, and merge at bottom in either *pcp1-AID* (MKSP3773) or *cmp7Δpcp1-AID* (MKSP3775) strains expressing OsTir1F74A. Cells were treated with DMSO or 5IAA as indicated. Scale bar, 1 µm. **C:** Plot of the percentage of *pcp1-AID* (gray bars, MKSP3773) or *cmp7Δpcp1-AID* (teal bars, MKSP3775) MGM4-expressing cells with N/C fluorescence ratios greater than 0.75 reflecting a loss of nuclear integrity. Points are the mean from each biological replicate, while bars and error bars represent total mean and standard deviation, respectively. Points represent average of at least 18 nuclei/cell cycle phase/strain/replicate, N = 649 nuclei. **D:** Western blot (WB) of Ppc89-AID levels (detected with an α-AID antibody) in total protein extracts from cells expressing OsTir1F74A and treated with 5IAA as indicated. Ponceau stain is shown at bottom to assess relative total protein loads. (-)OsTir1F74A (MKSP4175); (+)OsTir1F74A (MKSP4136); (-)OsTir1F74A, *cmp7Δ* (MKSP4176); (+)OsTir1F74A, *cmp7Δ* (MKSP4135). **E:** Fluorescence micrographs of representative nuclei are shown arranged vertically with MGM4 (green) at top, Cut11-mCherry (magenta) and mCherry-Atb2 (magenta) in middle, and merge at bottom in either *ppc89-AID* (MKSP4136) or *cmp7Δppc89-AID* (MKSP4135) strains expressing OsTir1F74A. Cells were treated with DMSO or 5IAA as indicated. Scale bar, 1 µm. **F:** Plot of the percentage of *ppc89-AID* (gray bars, MKSP4136) or *cmp7Δppc89-AID* (teal bars, MKSP4135) MGM4-expressing cells with N/C fluorescence ratios greater than 0.75 reflecting a loss of nuclear integrity. Points are the mean from each biological replicate, while bars and error bars represent total mean and standard deviation, respectively. Points represent average of at least 11 nuclei/cell cycle phase/strain/replicate, N = 504 nuclei. **G:** Time-lapse fluorescence microscopy of Atb2-mCherry and Cut11-mCherry in a mitotic *pcp1-AID* (MKSP3773) or *cmp7Δpcp1-AID* cell (MKSP3775) expressing OsTir1F74A. Imaging was begun after 1 h treatment with either DMSO or 5IAA as indicated. Scale bar, 1 µm. **H:** Time-lapse fluorescence microscopy of Cut11-mCherry in a mitotic *ppc89-AID* (MKSP4136) or *cmp7Δppc89-AID* cell (MKSP4135) expressing OsTir1F74A. Imaging was begun after 1 h treatment with either DMSO or 5IAA as indicated. Scale bar, 1 µm. **I:** Fluorescence micrographs of a *pcp1-AID* cell (MKSP3773) expressing OsTir1F74A containing a monopolar spindle arranged horizontally with MGM4 (green) at left, Cut11-mCherry (magenta) and mCherry-Atb2 (magenta) in middle, and merge on right 3 h after treatment with 5IAA. Scale bar, 1 µm.

**Supplementary Figure S6.**
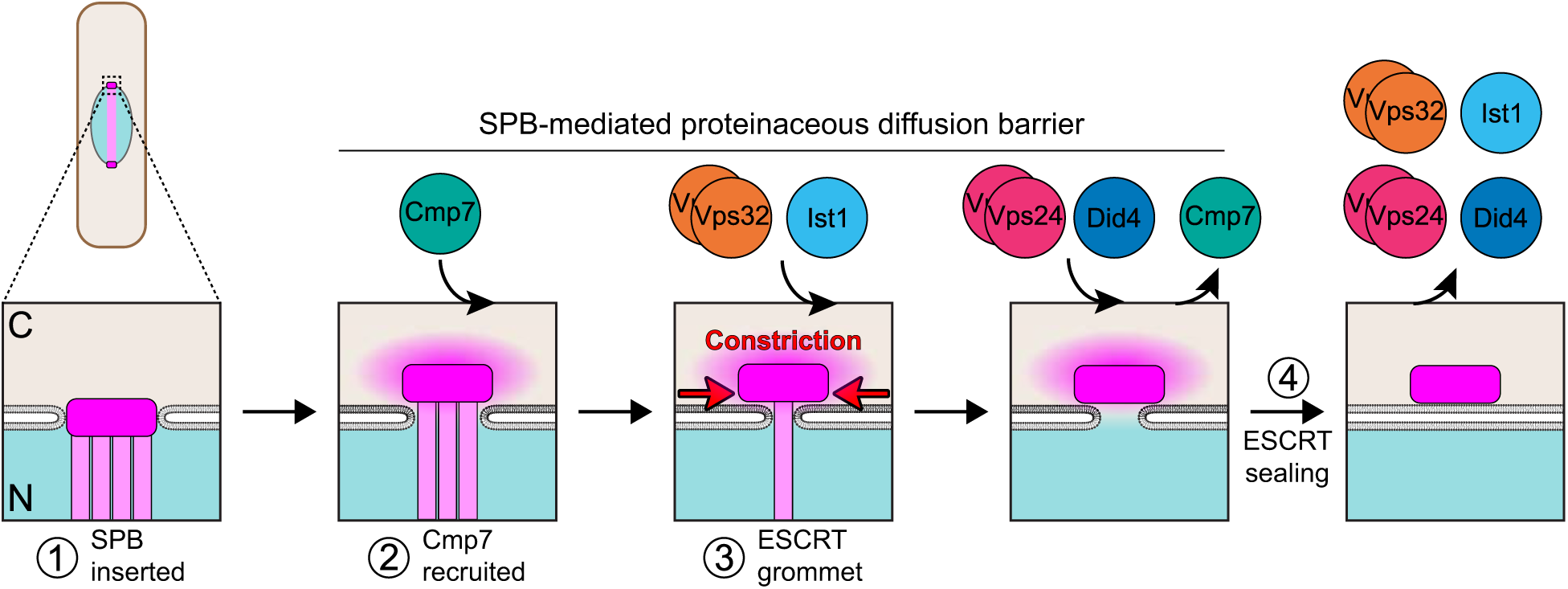
An ESCRT grommet and SPB-dependent diffusion barrier maintain nuclear integrity. A model of NE sealing and maintenance of mitotic nuclear integrity by the ESCRT machinery and the SPB. (1) In early mitosis, the SPB is inserted into the NE. (2) Once the SPB is extruded to the cytosol, Cmp7 is recruited by Heh1/Lem2 (not shown). Concurrently, the SPB contributes to the formation of a proteinaceous diffusion barrier sensitive to 2.5% 1,6-HD to prevent free diffusion through the unsealed NE. (3) A second wave of ESCRT recruitment comprising Vps32 and Ist1 occurs in late anaphase B, which generates a “grommet” that restricts the size of the NE hole and supports the SPB-dependent diffusion barrier. (4) Finally, in the G1/S of the following cell cycle, the third wave of ESCRTs, Did4 and Vps24, is recruited, displacing Cmp7 and sealing the NE.

## MOVIES

**Movie 1. CLEM and ET of a WT cell in early anaphase B.** Electron tomogram of an early anaphase B cell (MKSP3180) with segmentation. NE, blue; SPB, dark orange; microtubules, yellow. Scale bar, 100 nm.

**Movie 2. CLEM and ET of a WT cell in late anaphase B.** Electron tomogram of a late anaphase B cell (MKSP3180) with segmentation. NE, blue; SPB, dark orange; microtubules, yellow. Annotations at end of movie show NPC and SPB extrusion hole. Scale bar, 100 nm.

**Movie 3. CLEM and ET of a *cmp7Δ* cell in early anaphase B.** Electron tomogram of an early anaphase B cell (MKSP3243) with segmentation. NE, blue; SPB, dark orange; microtubules, yellow. Scale bar, 100 nm.

**Movie 4. CLEM and ET of a *cmp7Δ* cell in late anaphase B.** Electron tomogram of a late anaphase B cell (MKSP3243) with segmentation. NE, blue; SPB, dark orange; microtubules, yellow. Scale bar, 100 nm.

## SUPPLEMENTARY TABLES

**Supplementary Table 1.** GFP-tagging of key ESCRT machinery. ESCRT protein in *S. pombe* is shown on left. Orthologs in *S. cerevisiae* and *H. sapiens* are shown in center. Tagging approach to ensure function is shown on right.

| Protein | Orthologs |  | GFP tagging approach |
| --- | --- | --- | --- |
| *S. pombe* | *S. cerevisiae* | *H. sapiens* |  |
| Cmp7 | Chm7 | CHMP7 | Endogenous, C-terminal 3×HA-mEGFP |
| Vps32 | Snf7 | CHMP4A-C | Ectopic in *vps32Δ*, N-terminal 3×HA-mEGFP |
| Ist1 | Ist1 | IST1 | Endogenous, C-terminal 3×HA-mEGFP |
| Vps24 | Vps24 | CHMP3 | Ectopic in *vps24Δ*, N-terminal 3×HA-mEGFP |
| Did4 | Vps2 | CHMP2A,B | Ectopic in *did4Δ*, N-terminal 3×HA-mEGFP |
| Did2 | Did2 | CHMP1A,B | Endogenous, C-terminal 3×HA-mEGFP |

**Supplementary Table 2.**
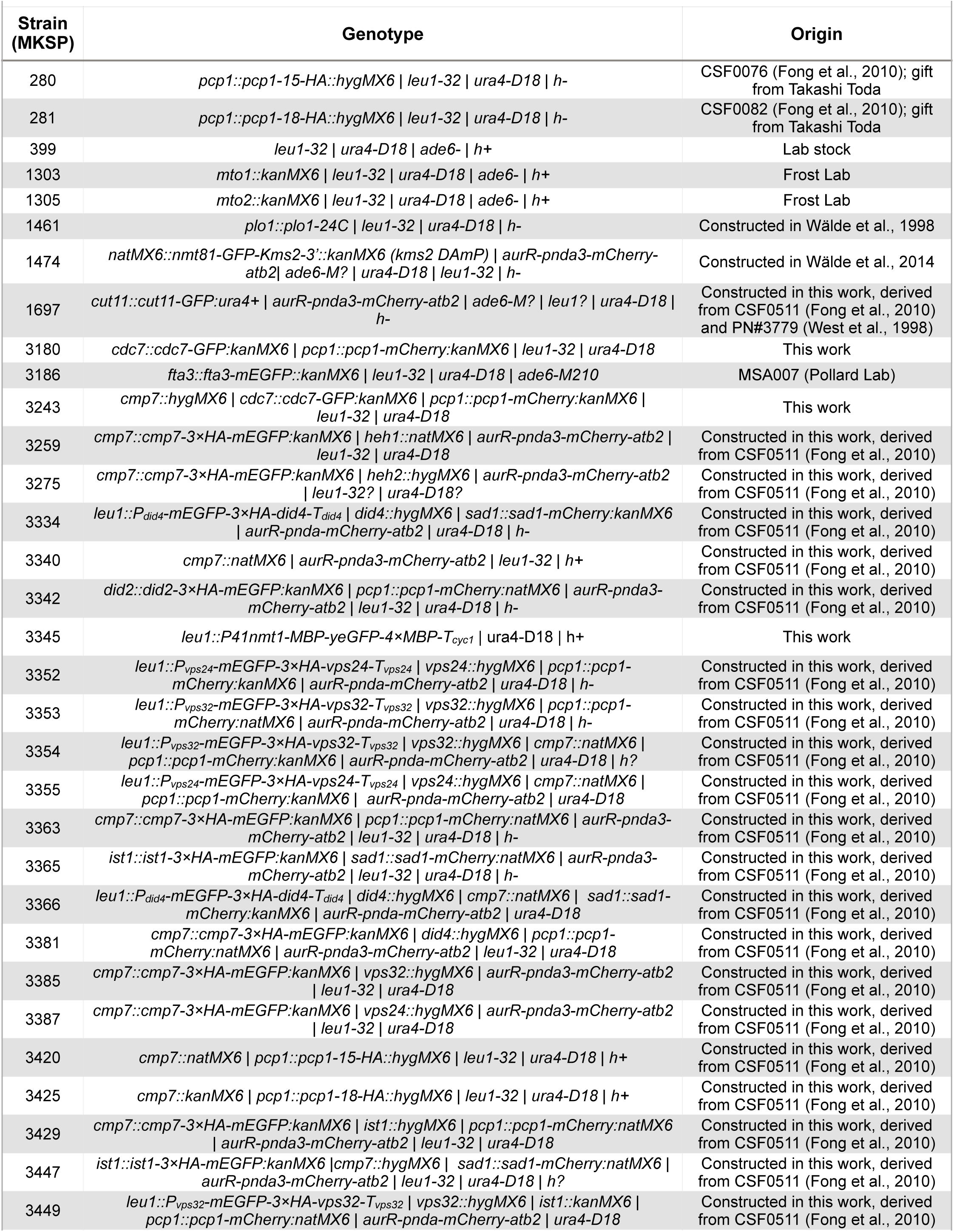
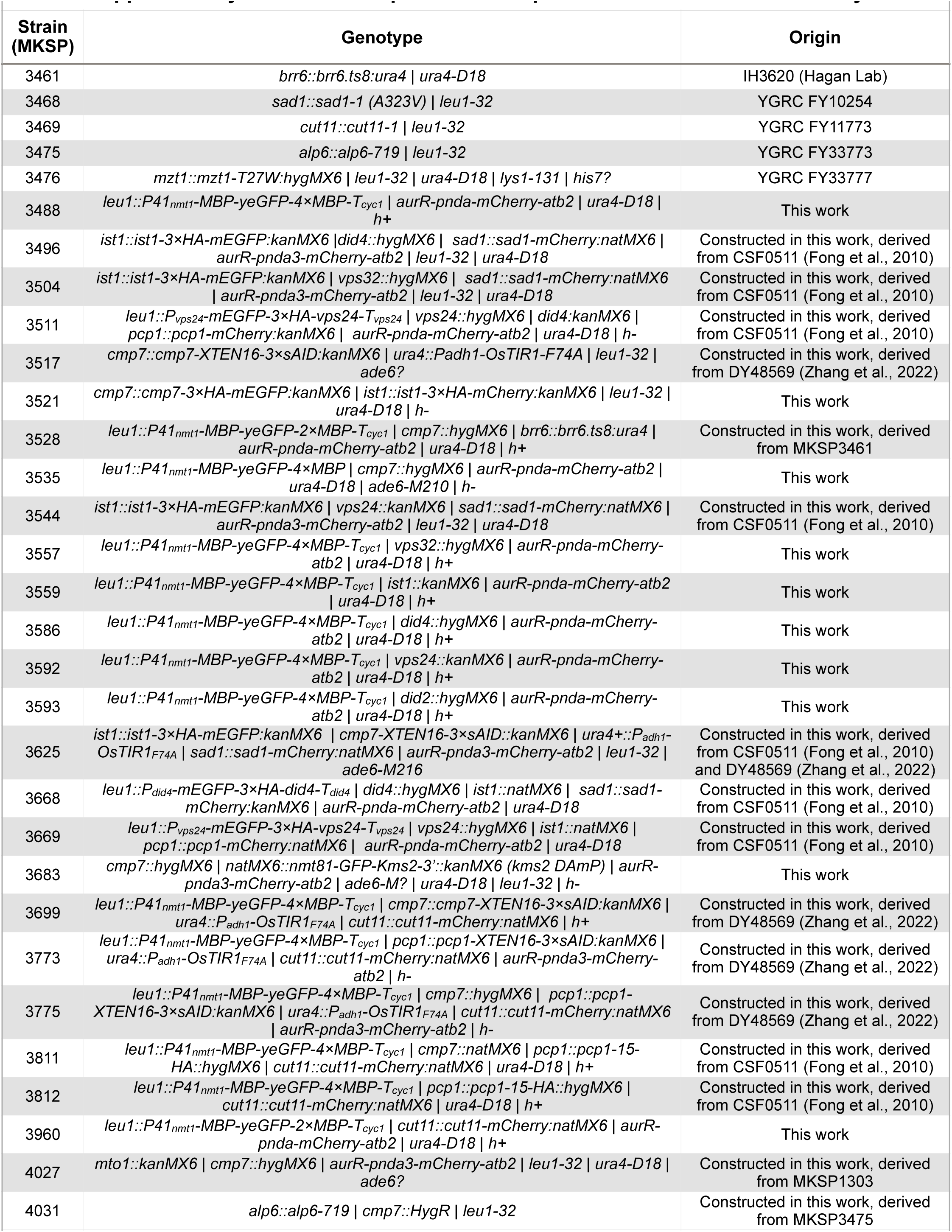
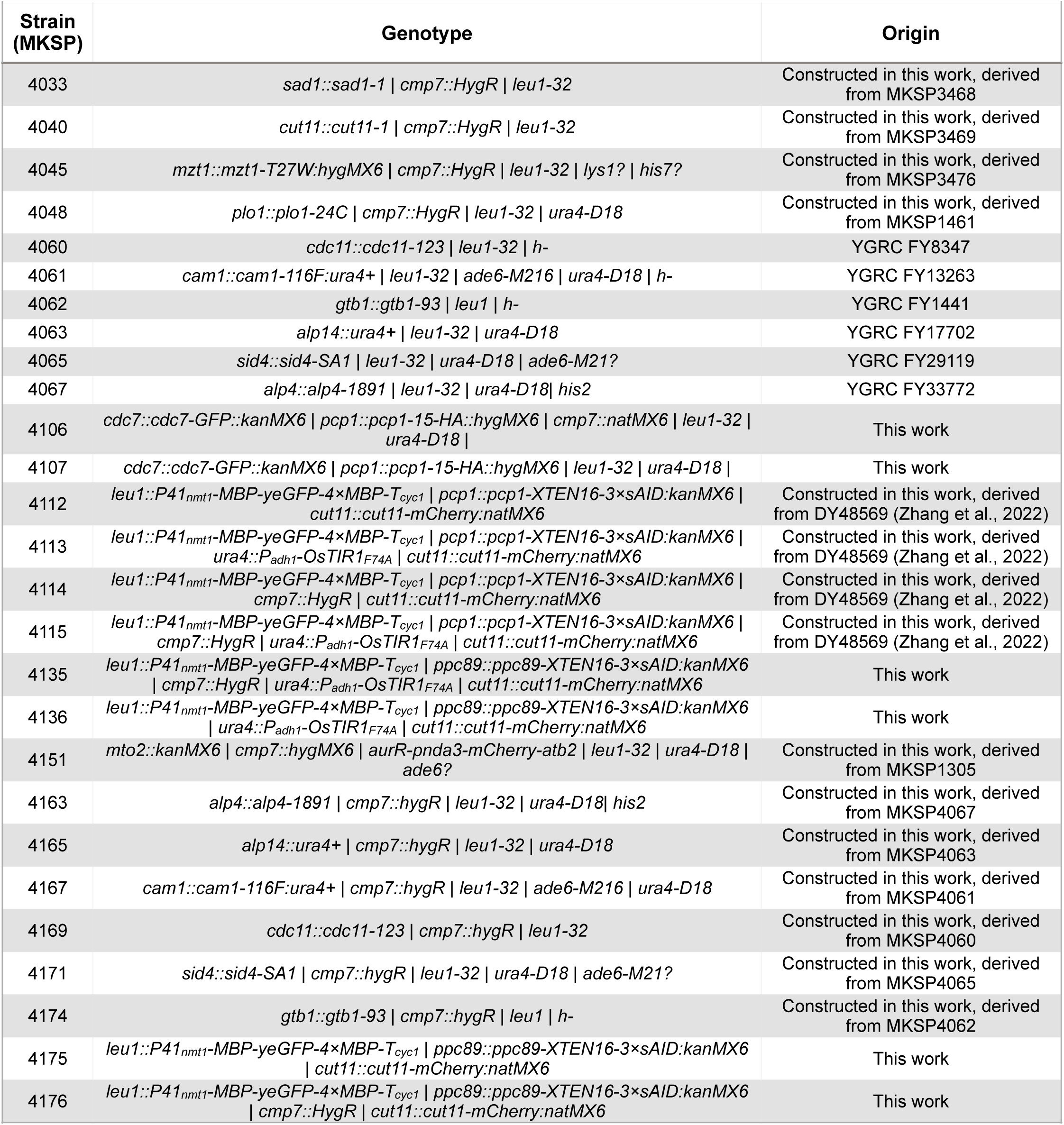
List of *S. pombe* strains used in this study.

**Supplementary Table 3.** List of plasmids used in this study.

| Identifier | Name/Description | Usage | Origin |
| --- | --- | --- | --- |
| pNA8 | pFA6a-3×HA-mEGFP-KanMX6 | Template for PCR based chromosomal integration of 3×HA-mEGFP ORF | This study; Addgene 87023 |
| pNA22 | pJK148-P<sub>vps32</sub>-mEGFP-3×HA-vps32-T<sub>vps32</sub> | Linearized with NruI and transformed into MKSP975 | This study, Gibson assembly |
| pNA23 | pJK148-P<sub>vps24</sub>-mEGFP-3×HA-vps24-T<sub>vps24</sub> | Linearized with NruI and transformed into MKSP975 | This study, Gibson assembly |
| pNA33 | pFA6a-hygMX6 | Integration cassette amplification hygMX6 template | Hentges et al., 2005 |
| pNA34 | pFA6a-natMX6 | Integration cassette amplification natMX6 template | Hentges et al., 2005 |
| pNA35 | pFA6a-kanMX6 | Template for PCR based chromosomal integration of kanMX6 ORF | Wach et al., 1997; Addgene 39296 |
| pNA45 | pJK148-P<sub>did4</sub>-mEGFP-3×HA-did4-T<sub>did4</sub> | Linearized with NruI and transformed into MKSP975 | This study, Gibson assembly |
| pNA47 | pFA6a-mCherry-KanMX6 | Template for PCR based chromosomal integration of mCherry ORF | EUROSCARF |
| pNA48 | pFA6a-mCherry-NatMX6 | Template for PCR based chromosomal integration of mCherry ORF | EUROSCARF |
| pNA50 | pFA6a-3×HA-mCherry-NatMX6 | Template for PCR based chromosomal integration of 3×HA-mCherry ORF | This study |
| pNA55 | pDB4581_pFA6a-3×sAID-KanMX6 | Template for PCR based chromosomal integration of 3×sAID ORF | Zhang et al., 2022; Addgene 171124 |
| pNA63 | pJK148-P41<sub>nmt1</sub>-MBP-yeGFP-4×MBP-T<sub>cyc1</sub> | Linearized with NruI and transformed into MKSP399 | Popken et al., 2015 and this study, restriction digest cloning |

## REFERENCES

1. Adell, M.A.Y., Migliano, S.M., Upadhyayula, S., Bykov, Y.S., Sprenger, S., Pakdel, M., Vogel, G.F., Jih, G., Skillern, W., Behrouzi, R., et al. (2017). Recruitment dynamics of ESCRT-III and Vps4 to endosomes and implications for reverse membrane budding. Elife 6, e31652. 10.7554/eLife.31652.

2. Babst, M., Katzmann, D.J., Estepa-Sabal, E.J., Meerloo, T., and Emr, S.D. (2002). ESCRT-III: an endosome-associated heterooligomeric protein complex required for MVB sorting. Dev Cell 3, 271–282. 10.1016/s1534-5807(02)00220-4.

3. Bahler, J., Wu, J.Q., Longtine, M.S., Shah, N.G., McKenzie, A., 3rd, Steever, A.B., Wach, A., Philippsen, P., and Pringle, J.R. (1998). Heterologous modules for efficient and versatile PCR-based gene targeting in Schizosaccharomyces pombe. Yeast 14, 943–951. 10.1002/(SICI)1097-0061(199807)14:10<943::AID-YEA292>3.0.CO;2-Y.

4. Byers, B., and Goetsch, L. (1975). Behavior of spindles and spindle plaques in the cell cycle and conjugation of Saccharomyces cerevisiae. J Bacteriol 124, 511–523. 10.1128/jb.124.1.511-523.1975.

5. Celetti, G., Paci, G., Caria, J., VanDelinder, V., Bachand, G., and Lemke, E.A. (2020). The liquid state of FG-nucleoporins mimics permeability barrier properties of nuclear pore complexes. The Journal of cell biology 219. 10.1083/jcb.201907157.

6. Chiu, Y.P., Sun, Y.C., Qiu, D.C., Lin, Y.H., Chen, Y.Q., Kuo, J.C., and Huang, J.R. (2020). Liquid-liquid phase separation and extracellular multivalent interactions in the tale of galectin-3. Nature communications 11, 1229. 10.1038/s41467-020-15007-3.

7. Coyne, A.N., Baskerville, V., Zaepfel, B.L., Dickson, D.W., Rigo, F., Bennett, F., Lusk, C.P., and Rothstein, J.D. (2021). Nuclear accumulation of CHMP7 initiates nuclear pore complex injury and subsequent TDP-43 dysfunction in sporadic and familial ALS. Sci Transl Med 13. 10.1126/scitranslmed.abe1923.

8. Davis, M.W., and Jorgensen, E.M. (2022). ApE, A Plasmid Editor: a freely available DNA manipulation and visualization program. Front Bioinform 2, 818619. 10.3389/fbinf.2022.818619.

9. Denais, C.M., Gilbert, R.M., Isermann, P., McGregor, A.L., te Lindert, M., Weigelin, B., Davidson, P.M., Friedl, P., Wolf, K., and Lammerding, J. (2016). Nuclear envelope rupture and repair during cancer cell migration. Science 352, 353–358. 10.1126/science.aad7297.

10. Dey, G., Culley, S., Curran, S., Schmidt, U., Henriques, R., Kukulski, W., and Baum, B. (2020). Closed mitosis requires local disassembly of the nuclear envelope. Nature 585, 119–123. 10.1038/s41586-020-2648-3.

11. Ding, R., West, R.R., Morphew, D.M., Oakley, B.R., and McIntosh, J.R. (1997). The spindle pole body of Schizosaccharomyces pombe enters and leaves the nuclear envelope as the cell cycle proceeds. Molecular biology of the cell 8, 1461–1479. 10.1091/mbc.8.8.1461.

12. Dultz, E., Wojtynek, M., Medalia, O., and Onischenko, E. (2022). The nuclear pore complex: birth, life, and death of a cellular behemoth. Cells 11. 10.3390/cells11091456.

13. Feng, Z., Caballe, A., Wainman, A., Johnson, S., Haensele, A.F.M., Cottee, M.A., Conduit, P.T., Lea, S.M., and Raff, J.W. (2017). Structural basis for mitotic centrosome assembly in flies. Cell 169, 1078–1089 e1013. 10.1016/j.cell.2017.05.030.

14. Flory, M.R., Morphew, M., Joseph, J.D., Means, A.R., and Davis, T.N. (2002). Pcp1p, an Spc110p-related calmodulin target at the centrosome of the fission yeast Schizosaccharomyces pombe. Cell Growth Differ 13, 47–58.

15. Fong, C.S., Sato, M., and Toda, T. (2010). Fission yeast Pcp1 links polo kinase-mediated mitotic entry to gamma-tubulin-dependent spindle formation. The EMBO journal 29, 120–130. 10.1038/emboj.2009.331.

16. Frey, S., Richter, R.P., and Görlich, D. (2006). FG-rich repeats of nuclear pore proteins form a three-dimensional meshwork with hydrogel-like properties. Science 314, 815–817. 10.1126/science.1132516.

17. Gibson, D.G., Young, L., Chuang, R.Y., Venter, J.C., Hutchison, C.A., 3rd, and Smith, H.O. (2009). Enzymatic assembly of DNA molecules up to several hundred kilobases. Nature methods 6, 343–345. 10.1038/nmeth.1318.

18. Gu, M., LaJoie, D., Chen, O.S., von Appen, A., Ladinsky, M.S., Redd, M.J., Nikolova, L., Bjorkman, P.J., Sundquist, W.I., Ullman, K.S., and Frost, A. (2017). LEM2 recruits CHMP7 for ESCRT-mediated nuclear envelope closure in fission yeast and human cells. Proceedings of the National Academy of Sciences of the United States of America 114, E2166–E2175. 10.1073/pnas.1613916114.

19. Halfmann, C.T., Sears, R.M., Katiyar, A., Busselman, B.W., Aman, L.K., Zhang, Q., O’Bryan, C.S., Angelini, T.E., Lele, T.P., and Roux, K.J. (2019). Repair of nuclear ruptures requires barrier-to-autointegration factor. The Journal of cell biology 218, 2136–2149. 10.1083/jcb.201901116.

20. Harris, M.A., Rutherford, K.M., Hayles, J., Lock, A., Bahler, J., Oliver, S.G., Mata, J., and Wood, V. (2022). Fission stories: using PomBase to understand Schizosaccharomyces pombe biology. Genetics 220. 10.1093/genetics/iyab222.

21. Henne, W.M., Buchkovich, N.J., Zhao, Y., and Emr, S.D. (2012). The endosomal sorting complex ESCRT-II mediates the assembly and architecture of ESCRT-III helices. Cell 151, 356–371. 10.1016/j.cell.2012.08.039.

22. Hentges, P., Van Driessche, B., Tafforeau, L., Vandenhaute, J., and Carr, A.M. (2005). Three novel antibiotic marker cassettes for gene disruption and marker switching in Schizosaccharomyces pombe. Yeast 22, 1013–1019. 10.1002/yea.1291.

23. Huxley, C., Green, E.D., and Dunham, I. (1990). Rapid assessment of S. cerevisiae mating type by PCR. Trends Genet 6, 236. 10.1016/0168-9525(90)90190-h.

24. Jaspersen, S.L. (2021). Anatomy of the fungal microtubule organizing center, the spindle pole body. Curr Opin Struct Biol 66, 22–31. 10.1016/j.sbi.2020.09.008.

25. Jia, J., Claude-Taupin, A., Gu, Y., Choi, S.W., Peters, R., Bissa, B., Mudd, M.H., Allers, L., Pallikkuth, S., Lidke, K.A., et al. (2020). Galectin-3 coordinates a cellular system for lysosomal repair and removal. Dev Cell 52, 69–87 e68. 10.1016/j.devcel.2019.10.025.

26. Jiang, X., Ho, D.B.T., Mahe, K., Mia, J., Sepulveda, G., Antkowiak, M., Jiang, L., Yamada, S., and Jao, L.E. (2021). Condensation of pericentrin proteins in human cells illuminates phase separation in centrosome assembly. Journal of cell science 134. 10.1242/jcs.258897.

27. Kanke, M., Nishimura, K., Kanemaki, M., Kakimoto, T., Takahashi, T.S., Nakagawa, T., and Masukata, H. (2011). Auxin-inducible protein depletion system in fission yeast. BMC Cell Biol 12, 8. 10.1186/1471-2121-12-8.

28. Keeney, J.B., and Boeke, J.D. (1994). Efficient targeted integration at leu1-32 and ura4-294 in Schizosaccharomyces pombe. Genetics 136, 849–856.

29. Kinugasa, Y., Hirano, Y., Sawai, M., Ohno, Y., Shindo, T., Asakawa, H., Chikashige, Y., Shibata, S., Kihara, A., Haraguchi, T., and Hiraoka, Y. (2019). The very-long-chain fatty acid elongase Elo2 rescues lethal defects associated with loss of the nuclear barrier function in fission yeast cells. Journal of cell science 132, jcs229021. 10.1242/jcs.229021.

30. Kono, Y., Adam, S.A., Sato, Y., Reddy, K.L., Zheng, Y., Medalia, O., Goldman, R.D., Kimura, H., and Shimi, T. (2022). Nucleoplasmic lamin C rapidly accumulates at sites of nuclear envelope rupture with BAF and cGAS. The Journal of cell biology 221, e202201024. 10.1083/jcb.202201024.

31. Kremer, J.R., Mastronarde, D.N., and McIntosh, J.R. (1996). Computer visualization of three-dimensional image data using IMOD. J Struct Biol 116, 71–76. 10.1006/jsbi.1996.0013.

32. Kroschwald, S., Maharana, S., and Simon, A. (2017). Hexanediol: a chemical probe to investigate the material properties of membrane-less compartments. Matters 3. 10.19185/matters.201702000010.

33. Krüger, L.K., Sanchez, J.L., Paoletti, A., and Tran, P.T. (2019). Kinesin-6 regulates cell-size-dependent spindle elongation velocity to keep mitosis duration constant in fission yeast. Elife 8, e42182. 10.7554/eLife.42182.

34. Kukulski, W., Schorb, M., Welsch, S., Picco, A., Kaksonen, M., and Briggs, J.A. (2012). Precise, correlated fluorescence microscopy and electron tomography of lowicryl sections using fluorescent fiducial markers. Methods Cell Biol 111, 235–257. 10.1016/B978-0-12-416026-2.00013-3.

35. Kume, K., Cantwell, H., Burrell, A., and Nurse, P. (2019). Nuclear membrane protein Lem2 regulates nuclear size through membrane flow. Nature communications 10, 1871. 10.1038/s41467-019-09623-x.

36. Lawrimore, J., Bloom, K.S., and Salmon, E.D. (2011). Point centromeres contain more than a single centromere-specific Cse4 (CENP-A) nucleosome. The Journal of cell biology 195, 573–582. 10.1083/jcb.201106036.

37. Lee, I.J., Stokasimov, E., Dempsey, N., Varberg, J.M., Jacob, E., Jaspersen, S.L., and Pellman, D. (2020). Factors promoting nuclear envelope assembly independent of the canonical ESCRT pathway. The Journal of cell biology 219. 10.1083/jcb.201908232.

38. Makarova, M., and Oliferenko, S. (2016). Mixing and matching nuclear envelope remodeling and spindle assembly strategies in the evolution of mitosis. Curr Opin Cell Biol 41, 43–50. 10.1016/j.ceb.2016.03.016.

39. Mastronarde, D.N. (2005). Automated electron microscope tomography using robust prediction of specimen movements. J Struct Biol 152, 36–51. 10.1016/j.jsb.2005.07.007.

40. Mastronarde, D.N., and Held, S.R. (2017). Automated tilt series alignment and tomographic reconstruction in IMOD. J Struct Biol 197, 102–113. 10.1016/j.jsb.2016.07.011.

41. McCullough, J., Clippinger, A.K., Talledge, N., Skowyra, M.L., Saunders, M.G., Naismith, T.V., Colf, L.A., Afonine, P., Arthur, C., Sundquist, W.I., et al. (2015). Structure and membrane remodeling activity of ESCRT-III helical polymers. Science 350, 1548–1551. 10.1126/science.aad8305.

42. McCullough, J., Frost, A., and Sundquist, W.I. (2018). Structures, functions, and dynamics of ESCRT-III/Vps4 membrane remodeling and fission complexes. Annu Rev Cell Dev Biol 34, 85–109. 10.1146/annurev-cellbio-100616-060600.

43. McDonald, K. (1999). High-pressure freezing for preservation of high resolution fine structure and antigenicity for immunolabeling. Methods Mol Biol 117, 77–97. 10.1385/1-59259-201-5:77.

44. Mierzwa, B.E., Chiaruttini, N., Redondo-Morata, L., von Filseck, J.M., Konig, J., Larios, J., Poser, I., Muller-Reichert, T., Scheuring, S., Roux, A., and Gerlich, D.W. (2017). Dynamic subunit turnover in ESCRT-III assemblies is regulated by Vps4 to mediate membrane remodelling during cytokinesis. Nature cell biology 19, 787–798. 10.1038/ncb3559.

45. Moreno, S., Klar, A., and Nurse, P. (1991). Molecular genetic analysis of fission yeast Schizosaccharomyces pombe. Methods Enzymol 194, 795–823. 10.1016/0076-6879(91)94059-l.

46. Murray, J.M., Watson, A.T., and Carr, A.M. (2016). Transformation of Schizosaccharomyces pombe: lithium acetate/dimethyl sulfoxide procedure. Cold Spring Harbor protocols 2016, pdb prot090969. 10.1101/pdb.prot090969.

47. Nguyen, H.C., Talledge, N., McCullough, J., Sharma, A., Moss, F.R., 3rd, Iwasa, J.H., Vershinin, M.D., Sundquist, W.I., and Frost, A. (2020). Membrane constriction and thinning by sequential ESCRT-III polymerization. Nat Struct Mol Biol 27, 392–399. 10.1038/s41594-020-0404-x.

48. O’Toole, E.T., Winey, M., and McIntosh, J.R. (1999). High-voltage electron tomography of spindle pole bodies and early mitotic spindles in the yeast Saccharomyces cerevisiae. Molecular biology of the cell 10, 2017–2031. 10.1091/mbc.10.6.2017.

49. Olmos, Y., Hodgson, L., Mantell, J., Verkade, P., and Carlton, J.G. (2015). ESCRT-III controls nuclear envelope reformation. Nature 522, 236–239. 10.1038/nature14503.

50. Olmos, Y., Perdrix-Rosell, A., and Carlton, J.G. (2016). Membrane binding by CHMP7 coordinates ESCRT-III-dependent nuclear envelope reformation. Current biology : CB 26, 2635–2641. 10.1016/j.cub.2016.07.039.

51. Penfield, L., Shankar, R., Szentgyorgyi, E., Laffitte, A., Mauro, M.S., Audhya, A., Muller-Reichert, T., and Bahmanyar, S. (2020). Regulated lipid synthesis and LEM2/CHMP7 jointly control nuclear envelope closure. The Journal of cell biology 219. 10.1083/jcb.201908179.

52. Petri, M., Frey, S., Menzel, A., Görlich, D., and Techert, S. (2012). Structural characterization of nanoscale meshworks within a nucleoporin FG hydrogel. Biomacromolecules 13, 1882–1889. 10.1021/bm300412q.

53. Pfitzner, A.K., Mercier, V., Jiang, X., Moser von Filseck, J., Baum, B., Saric, A., and Roux, A. (2020). An ESCRT-III polymerization sequence drives membrane deformation and fission. Cell 182, 1140–1155 e1118. 10.1016/j.cell.2020.07.021.

54. Pfitzner, A.K., Moser von Filseck, J., and Roux, A. (2021). Principles of membrane remodeling by dynamic ESCRT-III polymers. Trends Cell Biol 31, 856–868. 10.1016/j.tcb.2021.04.005.

55. Pieper, G.H., Sprenger, S., Teis, D., and Oliferenko, S. (2020). ESCRT-III/Vps4 controls heterochromatin-nuclear envelope attachments. Dev Cell 53, 27–41 e26. 10.1016/j.devcel.2020.01.028.

56. Popken, P., Ghavami, A., Onck, P.R., Poolman, B., and Veenhoff, L.M. (2015). Size-dependent leak of soluble and membrane proteins through the yeast nuclear pore complex. Molecular biology of the cell 26, 1386–1394. 10.1091/mbc.E14-07-1175.

57. Raab, M., Gentili, M., de Belly, H., Thiam, H.R., Vargas, P., Jimenez, A.J., Lautenschlaeger, F., Voituriez, R., Lennon-Dumenil, A.M., Manel, N., and Piel, M. (2016). ESCRT III repairs nuclear envelope ruptures during cell migration to limit DNA damage and cell death. Science 352, 359–362. 10.1126/science.aad7611.

58. Radulovic, M., Schink, K.O., Wenzel, E.M., Nahse, V., Bongiovanni, A., Lafont, F., and Stenmark, H. (2018). ESCRT-mediated lysosome repair precedes lysophagy and promotes cell survival. The EMBO journal 37. 10.15252/embj.201899753.

59. Remec Pavlin, M., and Hurley, J.H. (2020). The ESCRTs - converging on mechanism. Journal of cell science 133, jcs240333. 10.1242/jcs.240333.

60. Ribbeck, K., and Görlich, D. (2002). The permeability barrier of nuclear pore complexes appears to operate via hydrophobic exclusion. The EMBO journal 21, 2664–2671. 10.1093/emboj/21.11.2664.

61. Riquelme Barrientos, E., Otto, T.A., Mouton, S.N., Steen, A., and Veenhoff, L.M. (2023). Specificity and mechanism of 1,6 hexanediol-induced disruption of nuclear transport. bioRxiv, 2023.2003.2030.534880. 10.1101/2023.03.30.534880.

62. Rosenberg, J.A., Tomlin, G.C., McDonald, W.H., Snydsman, B.E., Muller, E.G., Yates, J.R., and Gould, K.L. (2006). Ppc89 links multiple proteins, including the septation initiation network, to the core of the fission yeast spindle-pole body. Molecular biology of the cell 17, 3793–3805. 10.1091/mbc.e06-01-0039.

63. Saksena, S., Wahlman, J., Teis, D., Johnson, A.E., and Emr, S.D. (2009). Functional reconstitution of ESCRT-III assembly and disassembly. Cell 136, 97–109. 10.1016/j.cell.2008.11.013.

64. Samwer, M., Schneider, M.W.G., Hoefler, R., Schmalhorst, P.S., Jude, J.G., Zuber, J., and Gerlich, D.W. (2017). DNA cross-bridging shapes a single nucleus from a set of mitotic chromosomes. Cell 170, 956–972 e923. 10.1016/j.cell.2017.07.038.

65. Schindelin, J., Arganda-Carreras, I., Frise, E., Kaynig, V., Longair, M., Pietzsch, T., Preibisch, S., Rueden, C., Saalfeld, S., Schmid, B., et al. (2012). Fiji: an open-source platform for biological-image analysis. Nature methods 9, 676–682. 10.1038/nmeth.2019.

66. Shulga, N., and Goldfarb, D.S. (2003). Binding dynamics of structural nucleoporins govern nuclear pore complex permeability and may mediate channel gating. Molecular and cellular biology 23, 534–542. 10.1128/MCB.23.2.534-542.2003.

67. Skibinski, G., Parkinson, N.J., Brown, J.M., Chakrabarti, L., Lloyd, S.L., Hummerich, H., Nielsen, J.E., Hodges, J.R., Spillantini, M.G., Thusgaard, T., et al. (2005). Mutations in the endosomal ESCRTIII-complex subunit CHMP2B in frontotemporal dementia. Nature genetics 37, 806–808. 10.1038/ng1609.

68. Skowyra, M.L., Schlesinger, P.H., Naismith, T.V., and Hanson, P.I. (2018). Triggered recruitment of ESCRT machinery promotes endolysosomal repair. Science 360, eaar5078. 10.1126/science.aar5078.

69. Sohrmann, M., Schmidt, S., Hagan, I., and Simanis, V. (1998). Asymmetric segregation on spindle poles of the Schizosaccharomyces pombe septum-inducing protein kinase Cdc7p. Genes & development 12, 84–94. 10.1101/gad.12.1.84.

70. Tamm, T., Grallert, A., Grossman, E.P., Alvarez-Tabares, I., Stevens, F.E., and Hagan, I.M. (2011). Brr6 drives the Schizosaccharomyces pombe spindle pole body nuclear envelope insertion/extrusion cycle. The Journal of cell biology 195, 467–484. 10.1083/jcb.201106076.

71. Teis, D., Saksena, S., and Emr, S.D. (2008). Ordered assembly of the ESCRT-III complex on endosomes is required to sequester cargo during MVB formation. Dev Cell 15, 578–589. 10.1016/j.devcel.2008.08.013.

72. Thaller, D.J., Allegretti, M., Borah, S., Ronchi, P., Beck, M., and Lusk, C.P. (2019). An ESCRT-LEM protein surveillance system is poised to directly monitor the nuclear envelope and nuclear transport system. Elife 8, e45284. 10.7554/eLife.45284.

73. Thaller, D.J., Tong, D., Marklew, C.J., Ader, N.R., Mannino, P.J., Borah, S., King, M.C., Ciani, B., and Lusk, C.P. (2021). Direct binding of ESCRT protein Chm7 to phosphatidic acid-rich membranes at nuclear envelope herniations. The Journal of cell biology 220. 10.1083/jcb.202004222.

74. Ungricht, R., and Kutay, U. (2017). Mechanisms and functions of nuclear envelope remodelling. Nature reviews. Molecular cell biology 18, 229–245. 10.1038/nrm.2016.153.

75. Ventimiglia, L.N., Cuesta-Geijo, M.A., Martinelli, N., Caballe, A., Macheboeuf, P., Miguet, N., Parnham, I.M., Olmos, Y., Carlton, J.G., Weissenhorn, W., and Martin-Serrano, J. (2018). CC2D1B coordinates ESCRT-III activity during the mitotic reformation of the nuclear envelope. Dev Cell 47, 547–563 e546. 10.1016/j.devcel.2018.11.012.

76. Vietri, M., Radulovic, M., and Stenmark, H. (2020). The many functions of ESCRTs. Nature reviews. Molecular cell biology 21, 25–42. 10.1038/s41580-019-0177-4.

77. Vietri, M., Schink, K.O., Campsteijn, C., Wegner, C.S., Schultz, S.W., Christ, L., Thoresen, S.B., Brech, A., Raiborg, C., and Stenmark, H. (2015). Spastin and ESCRT-III coordinate mitotic spindle disassembly and nuclear envelope sealing. Nature 522, 231–235. 10.1038/nature14408.

78. von Appen, A., LaJoie, D., Johnson, I.E., Trnka, M.J., Pick, S.M., Burlingame, A.L., Ullman, K.S., and Frost, A. (2020). LEM2 phase separation promotes ESCRT-mediated nuclear envelope reformation. Nature 582, 115–118. 10.1038/s41586-020-2232-x.

79. Wälde, S., and King, M.C. (2014). The KASH protein Kms2 coordinates mitotic remodeling of the spindle pole body. Journal of cell science 127, 3625–3640. 10.1242/jcs.154997.

80. Wallis, S.S., Ventimiglia, L.N., Otigbah, E., Infante, E., Cuesta-Geijo, M.A., Kidiyoor, G.R., Carbajal, M.A., Fleck, R.A., Foiani, M., Garcia-Manyes, S., et al. (2021). The ESCRT machinery counteracts Nesprin-2G-mediated mechanical forces during nuclear envelope repair. Dev Cell 56, 3192–3202 e3198. 10.1016/j.devcel.2021.10.022.

81. Webster, B.M., Colombi, P., Jäger, J., and Lusk, C.P. (2014). Surveillance of nuclear pore complex assembly by ESCRT-III/Vps4. Cell 159, 388–401. 10.1016/j.cell.2014.09.012.

82. Webster, B.M., Thaller, D.J., Jäger, J., Ochmann, S.E., Borah, S., and Lusk, C.P. (2016). Chm7 and Heh1 collaborate to link nuclear pore complex quality control with nuclear envelope sealing. The EMBO journal 35, 2447–2467. 10.15252/embj.201694574.

83. West, R.R., Vaisberg, E.V., Ding, R., Nurse, P., and McIntosh, J.R. (1998). cut11(+): A gene required for cell cycle-dependent spindle pole body anchoring in the nuclear envelope and bipolar spindle formation in Schizosaccharomyces pombe. Molecular biology of the cell 9, 2839–2855. 10.1091/mbc.9.10.2839.

84. Wood, V., Gwilliam, R., Rajandream, M.A., Lyne, M., Lyne, R., Stewart, A., Sgouros, J., Peat, N., Hayles, J., Baker, S., et al. (2002). The genome sequence of Schizosaccharomyces pombe. Nature 415, 871–880. 10.1038/nature724.

85. Woodruff, J.B., Ferreira Gomes, B., Widlund, P.O., Mahamid, J., Honigmann, A., and Hyman, A.A. (2017). The centrosome Is a selective condensate that nucleates microtubules by concentrating tubulin. Cell 169, 1066–1077 e1010. 10.1016/j.cell.2017.05.028.

86. Zhang, X.R., Zhao, L., Suo, F., Gao, Y., Wu, Q., Qi, X., and Du, L.L. (2022). An improved auxin-inducible degron system for fission yeast. G3 (Bethesda) 12, jkab393. 10.1093/g3journal/jkab393.

87. Zwicker, D., Decker, M., Jaensch, S., Hyman, A.A., and Julicher, F. (2014). Centrosomes are autocatalytic droplets of pericentriolar material organized by centrioles. Proceedings of the National Academy of Sciences of the United States of America 111, E2636–2645. 10.1073/pnas.1404855111.

